# Systematic comparison of pathogenic variants of the RNA exosome gene *EXOSC3/RRP40* in *Saccharomyces cerevisiae* reveals variant-specific functional consequences

**DOI:** 10.64898/2026.07.29.741578

**Authors:** Milo B. Fasken, Sara W. Leung, Leonardo F. Serafim, Chunli Yan, Madison L. Intemann, Dustin L. Gable, Kristin W. Barañano, Ivaylo Ivanov, Homa Ghalei, Anita H. Corbett

## Abstract

The RNA exosome is an essential, evolutionarily conserved ribonuclease complex that processes and degrades many classes of RNA. The complex is composed of three structural ‘cap’ subunits (EXOSC1-3/Csl4, Rrp4, Rrp40; *H. sapiens/S. cerevisiae*), six structural ‘core’ subunits (EXOSC4-9/Rrp41,Rrp46,Mtr3,Rrp42,Rrp43,Rrp45), and a catalytic ribonuclease (DIS3 or DIS3L/Dis3). Cofactors that associate with the RNA exosome confer specificity to target specific RNAs for processing and/or decay. Missense mutations in genes encoding structural subunits of the RNA exosome have been linked to neurological diseases. Notably, several pathogenic mutations have been identified in *EXOSC3* that are associated with pontocerebellar hypoplasia type 1b (PCH1b). These pathogenic alleles cause a broad spectrum of clinical severity, suggesting variant-specific functional consequences. Given the high degree of conservation between the human and budding yeast RNA exosome complexes, we performed a systematic analysis of eight pathogenic EXOSC3 variants modeled in budding yeast Rrp40. We find that two Rrp40 variants cause growth defects, show distinct negative genetic interactions with RNA exosome cofactor mutants, and impair RNA processing in budding yeast. One of these variants, EXOSC3-Y109N/Rrp40-Y64N, had not been previously characterized in any mechanistic studies. Computational stability predictions and immunoblot analyses indicate that most EXOSC3/Rrp40 variants display reduced steady-state protein levels, but decreased protein levels do not strictly correlate with phenotype or disease severity, suggesting that individual variants disrupt RNA exosome function through distinct mechanisms. Collectively, our studies suggest that pathogenic EXOSC3 variants alter RNA exosome function through distinct mechanisms and provide insight into the specific molecular defects that could underlie pathology.

## INTRODUCTION

The RNA exosome is an evolutionarily conserved, ubiquitously expressed complex (1) composed of nine structural subunits and a catalytic ribonuclease (2,3). The complex is required for essential processes such as the production of mature rRNAs in the nucleus, normal turnover of mRNAs in the cytoplasm, and degradation of aberrant RNAs in surveillance pathways in both the cytoplasm and the nucleus (4). The complex consists of three cap subunits (EXOSC1-3 in *H. sapiens*; Csl4,Rrp4,Rrp40 in *S. cerevisiae*) and six PH-like ring subunits (EXOSC4-9 in *H. sapiens*; Rrp41,Rrp46,Mtr3,Rrp42,Rrp43,Rrp45 in *S. cerevisiae*) that form a barrel-like core structure (5) (**Fig. 1A**). The structural ring subunits interact with a catalytic 3’-5’ riboexonuclease/endonuclease subunit (DIS3/DIS3L in *H. sapiens*; Dis3/Rrp44 in *S. cerevisiae*) at the bottom of the complex (6). In addition, the structural subunits interact with another 3’-5’ riboexonuclease (EXOSC10 in *H. sapiens*; Rrp6 in *S. cerevisiae*) at the top of the complex (7,8). Both the individual subunits and the overall structure of this essential complex are evolutionarily conserved (9–11).

**Figure 1.**
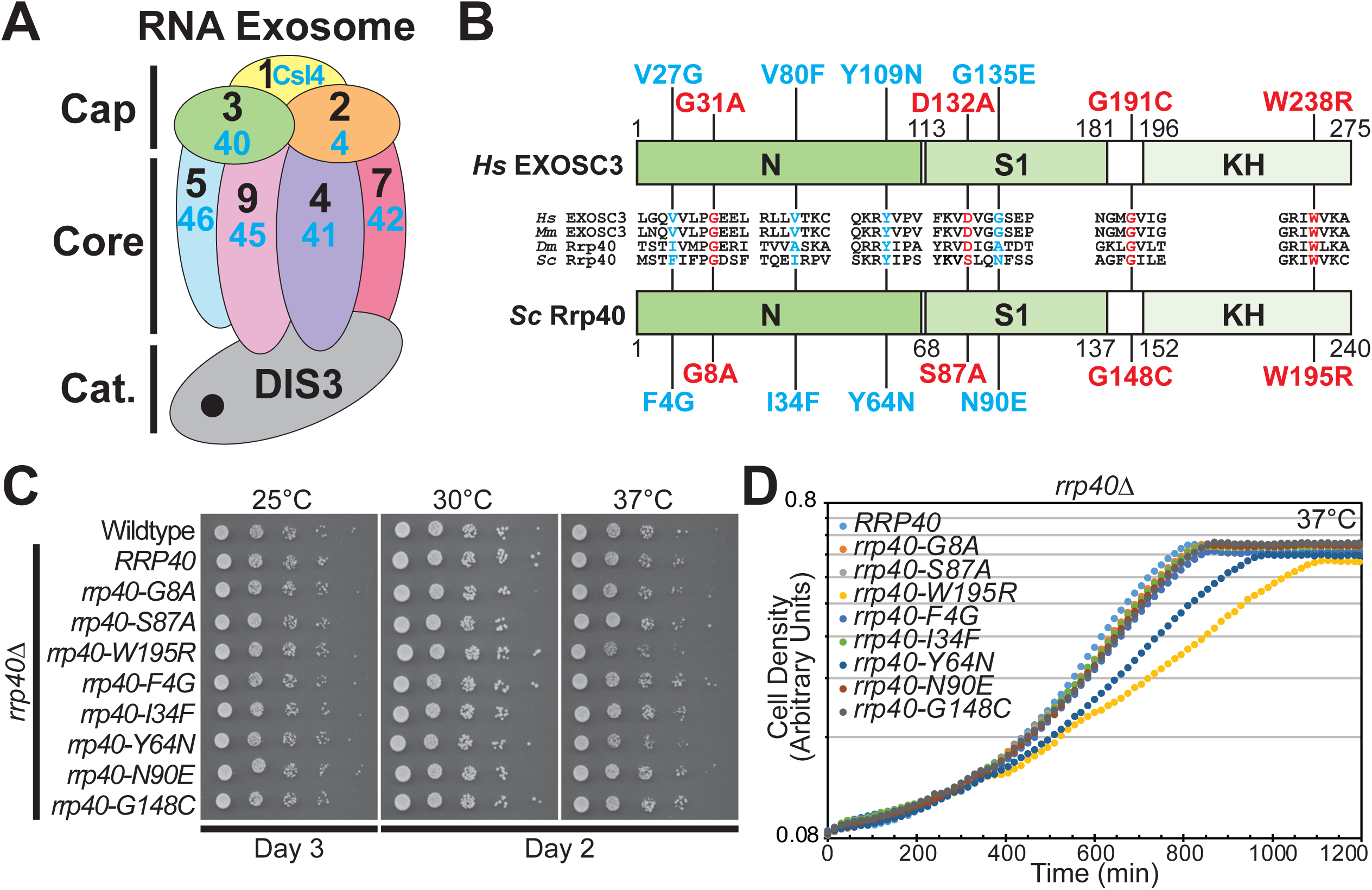
A subset of pathogenic EXOSC3 variants modeled in budding yeast Rrp40 impair cell growth. *A*, A cartoon model of the 10-subunit RNA exosome complex is shown. The subunits of the complex are indicated with both the human and budding yeast names indicated: EXOSC1-3/Csl1,Rrp4,Rrp40 in the cap and EXOSC4-9/Rrp41, Rrp46, Mtr3, Rrp42, Rrp43, Rrp45 in the barrel-like core ring, with the catalytic DIS3 subunit at the base. *B*, Schematic comparing domain organization of human EXOSC3 (*Hs* EXOSC3) and budding yeast Rrp40 (*Sc* Rrp40). EXOSC3/Rrp40 contains three domains: an N-terminal domain, a central S1 putative RNA binding domain, and a C-terminal putative RNA binding KH (K homology) domain. Above (EXOSC3) and below (Rrp40) are shown the pathogenic variants in EXOSC3 and corresponding modeled amino acid changes in Rrp40. Those amino acids indicated in red have been modeled in budding yeast previously (51,52) and those in blue have not. Alignment of EXOSC3/Rrp40 amino acid sequences from *Homo sapiens* (*Hs*), *Mus musculus* (*Mm*), *Drosophila melanogaster* (*Dm*), and *Saccharomyces cerevisiae* (*Sc*) show the conservation of the sequences immediately surrounding and the specific amino acids that are analyzed in this study. *C*,*D*, The *rrp40-W195R* and *rrp40-Y64N* mutant cells show impaired growth as analyzed by either a serial dilution growth assay (*C*) or growth in liquid media (*D*). *C*, WT control cells or *rrp40*Δ cells containing *RRP40* as a control or each *rpp40* mutant generated through a plasmid shuffle as described in Experimental Procedures indicated were serially diluted, spotted, and grown at 25, 30, or 37°C for 2-3 days as indicated. *D*, The same samples described in (*C*) were grown in liquid culture at 37°C for the time indicated. Results shown for cell growth are representative of at least three independent experiments.

The RNA exosome activity was initially identified in budding yeast in a screen for ribosomal RNA processing mutants (12), while biochemical purification defined the complex (13,14). Studies employing a variety of different approaches have defined many RNA targets of the RNA exosome including both coding and non-coding RNAs (15–19). Specificity for these target RNAs is achieved through various cofactors that associate with the RNA exosome in both the nucleus and the cytoplasm (20–22). Many of these evolutionarily conserved RNA exosome cofactors were initially identified in budding yeast, including the nuclear cofactors Mpp6 (23), Rrp47 (24), the RNA helicase Mtr4 (25), and the TRAMP complex, which contains Mtr4 (26,27). Other cofactors, primarily components of the SKI complex, associate with the RNA exosome in the cytoplasm (11,28,29). Recent work has identified several complexes that associate with the mammalian RNA exosome to confer target specificity, including the NEXT and PAXT complexes (19,22,30,31).

In recent years, a number of pathogenic missense variants have been identified in genes that encode structural subunits of the RNA exosome (9). Collectively, these diseases have been termed RNA exosomopathies. An initial report identified mutations in the *EXOSC3* gene in patients diagnosed with pontocerebellar hypoplasia type 1B (PCH1b) (32). Subsequent studies have identified additional pathogenic variants in *EXOSC3* as well as in *EXOSC1*, *8*, and *9* that have been linked to PCH1 (33–37). Mutations in *EXOSC2*, *4*, *5*, and *7* have also been linked to autosomal recessive diseases with brain defects (38–43). The majority of the pathogenic variants identified in *EXOSC* genes are missense mutations that alter single amino acids. The clinical presentations associated with these different *EXOSC* variants are diverse (9); however, many of the variants impact the cerebellum at least to some extent, consistent with the link to pontocerebellar hypoplasia. How variants that impact one ubiquitously expressed, essential complex cause distinct pathology is not yet clear.

PCH associated with mutations in the *EXOSC3* gene is characterized by congenital hypoplasia and/or progressive atrophy of the pons, the middle segment of the brainstem, and cerebellum (32,44,45). These structures contain pontocerebellar tracts critical for motor coordination, motor learning, and cognitive processing. PCH may also be associated with rare forms of spinal muscular atrophy, a lower motor neuron disorder caused by the loss and degeneration of alpha motor neurons within the spinal cord and brainstem which innervate skeletal muscles and enable voluntary movement (32,46). Together, pontocerebellar and motor neuron deficits produce a wide range of severe clinical manifestations arising from deficits of both the central and peripheral nervous systems. Affected patients often present at birth with profound hypotonia, microcephaly, and poor feeding and swallowing which may progress to respiratory failure and death in the neonatal or infantile period. In those who survive beyond the first year of life, spasticity, weakness, dystonia, and seizures have been reported (45,46). Life expectancy ranges from a few weeks in severe cases to a normal life span in mild cases, illustrating the diversity of clinical presentations even for mutations in a single gene (45).

Studies seeking to understand how pathogenic missense variants in genes encoding structural subunits of the RNA exosome impair the function the complex and contribute to pathology have employed a variety of different approaches. A number of studies have analyzed cells derived from patients, primarily fibroblasts (35,36,38,47). Other studies have employed genetic model systems, including budding yeast, zebrafish, and *Drosophila* (33,41,48–53) as well as engineered cell lines and computational approaches (54,55). A recent study extended these approaches by using CRISPR/Cas9 engineered human pluripotent stem cell derived cerebellar organoids (56). Results from these studies reveal that pathogenic RNA exosome subunit variants typically do impair RNA exosome function, but the mechanisms underlying these defects may be diverse.

Some studies demonstrate that the RNA exosomopathy variants cause a decrease in the steady-state level of both the directly affected subunit and other associated subunits (36,48), which could suggest an overall decrease in the level of the RNA exosome complex. However, the diversity of pathology observed in patients suggests that simple downregulation of the total complex levels is not sufficient to explain disease mechanism. Indeed, evidence suggests that the specific amino acid substitution may alter the function or interactions of the affected subunits (41,48). A recent study investigating missense variants in the S1 domain of EXOSC3 also demonstrated that these variants can destabilize the protein, weaken interactions with other RNA exosome cap subunits, and compromise RNA exosome complex integrity, leading to molecular defects, some of which can be rescued by increased levels of EXOSC3 (55). A number of the cofactors that facilitate the processing and degradation of specific RNA targets associate with the cap subunits of the RNA exosome, including EXOSC3 (22,57). Thus, changes in surface-exposed residues on EXOSC3 have the potential to impact key interactions with RNA exosome cofactors.

Since the initial report of pathogenic mutations in *EXOSC3* (32), a number of additional pathogenic mutations in *EXOSC3* have been identified with differing clinical severity (46,58–61). As a specific example, the initial report included one individual homozygous for c.92G>C (p.G31A) mutation (32). More individuals homozygous for this mutation have now been identified (60,62,63) and this specific mutation has been described as a founder mutation with high prevalence among the Roma population in certain regions (60). Another study reported two siblings who are compound heterozygous for variants in *EXOSC3* c.155delC and c.80T>G (p.V27G) with extraordinarily mild presentations of PCH1b (58). In these siblings, the first variant, p.P52RfsX2 (c.155delC) has been previously characterized as pathogenic (44), so the c.80T>G (p.V27G) was classified as a likely pathogenic variant. The V27 residue and the G31 residue are located close to one another both within the primary sequence of EXOSC3 and in the three-dimensional structure, raising the question of why such proximal changes confer such drastically different clinical outcomes. While genetic modifiers among affected individuals are likely to contribute to phenotypic diversity, another important factor may be the distinct ways in which specific amino acid changes impact RNA exosome function.

Here, we perform a systematic analysis modeling eight missense mutations identified in the *EXOSC3* gene (**Fig. 1B, Table S1**), including four reported pathogenic alleles that have not yet been analyzed at the molecular level, V27G (58), V80F (64), Y109N (45), and G135E (45). We employ a dual approach of modeling the changes in the budding yeast EXOSC3 orthologue encoded by *RRP40* and employing a cultured mammalian neuronal cell line. This strategy allows us to directly assess and compare the functional consequences of several reported pathogenic variants of EXOSC3. In addition, leveraging available RNA exosome complex structures and computational modeling, we provide protein stability analyses for each variant in both yeast and mammalian systems. This systematic analysis reveals that a model of the EXOSC3 Y109N variant in budding yeast causes slow growth and rRNA processing defect, providing the first molecular characterization of this variant. We also identify genetic interactions with specific RNA exosome cofactors, most notably Mpp6, which highlight the allele-specific defects caused by each mutation. An analysis of EXOSC3 steady-state levels and protein stability predictions show that many of the pathogenic variants likely alter the stability of the EXOSC3 protein, but these changes in protein levels do not correlate with disease severity, suggesting that a decrease in protein or complex levels is not the sole basis underlying disease. These findings highlight the importance of defining the functional consequences of specific pathogenic variants to help provide insight into the disease mechanism and to consider potential therapeutic interventions.

## Results

### Functional characterization of *EXOSC3/RRP40* missense variants using budding yeast model

To systematically model and compare a set of EXOSC3 variants that have been linked to disease, we modeled the corresponding amino acid changes in the budding yeast orthologue of EXOSC3, Rrp40 (**Fig. 1A,B**). While some of these variants have been previously modeled and analyzed in budding yeast (indicated in red in **Fig. 1B**) (50–52), others have not yet been modeled in this organism (indicated in blue in **Fig. 1B**). The patient genotypes and the clinical presentation of the variants shown in **Figure 1B**, are listed in **Table S1**. All of the modeled variants are derived from patients with PCH1b (46) and/or complicated spastic paraplegia (CSP), which is a group of inherited neurologic disorders marked by progressive lower-limb spasticity and weakness as well as additional neurologic features (65,66).

We modeled each of the pathogenic amino acid changes identified in patients in budding yeast, based on an alignment of human EXOSC3 with *S. cerevisiae* Rrp40 (**Fig. 1B**). To assess how each amino acid change impacts the function of the essential Rrp40 protein, we used a plasmid shuffle approach (67) to express each Rrp40 variant in budding yeast cells deleted for the endogenous *RRP40* gene (*rrp40*Δ). Cell growth was assessed by a serial dilution growth assay (**Fig. 1C**) as well as by growth in liquid media (**Fig. 1D**). The *rrp40*Δ cells transformed with control wildtype *RRP40* show growth at all temperatures tested (25°C, 30°C, 37°C) that is indistinguishable from growth of Wildtype control. The majority of the variants modeled do not impair growth. However, both *rrp40-W195R*, which models EXOSC3 W238R, and *rrp40-Y64N*, which models EXOSC3 Y109N, show slow growth at 37°C. This result is expected for *rrp40-W195R*, which has been studied previously (50–52), so this variant serves as a control for comparison to the previously uncharacterized *rrp40-Y64N* variant. Overall, these data suggest that many of the pathogenic variants present in patients do not severely impact the essential function of the RNA exosome in the budding yeast model. These results likely reflect the fact that the RNA exosome is essential for viability, even in budding yeast, such that variants causing severe functional impairment would not be compatible with viability, and, by extension, human development. Consistent with this idea, neither EXOSC3 W238R nor EXOSC3 Y109N has been reported as homozygous in patients (**Table S1**), suggesting that these variants may not support sufficient RNA exosome activity to permit normal human development.

### *rrp40* alleles that model disease variants exhibit negative genetic interactions with RNA exosome cofactors

To extend the functional analysis of the Rrp40 variants, we tested for genetic interactions between each *rrp40* mutant and deletion mutants of two, nonessential RNA exosome cofactors, Mpp6 and Rrp47, and deletion mutant of a nonessential RNA exosome catalytic subunit, Rrp6 (23,68,69). Mpp6, Rrp47 and Rrp6 interact with the cap subunits of the complex (20). As shown in the **Figure 2**, we analyzed the growth of the *rrp40 mpp6*Δ, *rrp40 rrp47*Δ, and *rrp40 rrp6*Δ double mutant cells by serial dilution in a solid media growth assay at both 30°C and 37°C (**Fig. 2A,C,E**) or by liquid growth assay at 37°C (**Fig. 2B,D,F**). Using this approach, we find that *rrp40-Y64N mpp6*Δ cells grow more slowly than *RRP40* Δ*mpp6* cells (**Fig. 2A,B**), which do not exhibit temperature-sensitive growth as previously observed (70), or *rrp40-Y64N* cells at 37°C (**Fig. 1C,D**). This result uncovers a negative genetic interaction between *rrp40-Y64N* and *mpp6*Δ. Notably, *rrp40-Y64N mpp6*Δ cells also grow more slowly than the *rrp40-W195R* Δ*mpp6* cells (**Fig. 2A,B**), suggesting the *mpp6*Δ deletion exerts a greater effect on the *rrp40-Y64N* mutant than on the *rrp40-W195R* mutant.

**Figure 2.**
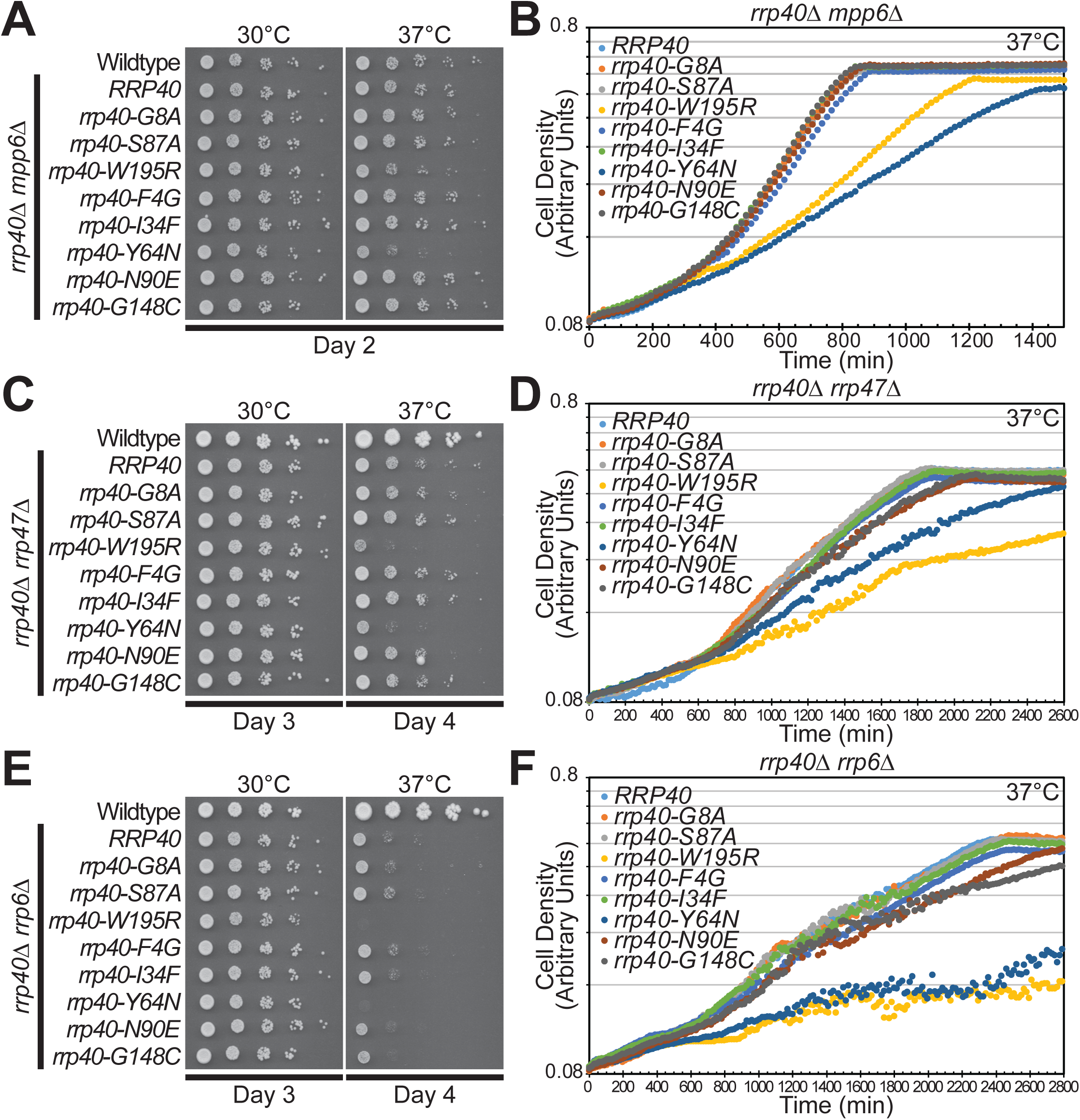
***rrp40-W195R* and *rrp40-Y64N* show genetic interactions with RNA exosome cofactors.** Each of the rrp40 models was combined with loss of a non-essential RNA exosome cofactor: Mpp6 (*A,B*), Rrp47 (*C,D*) or Rrp6 (*E,F*) and growth was analyzed by either serial dilution growth assay (*A,C,E*) or growth in liquid media (*B,D,F*). *A,* As described in Experimental Procedures, a plasmid shuffle approach was used in double mutant cells to examine each rrp40 variant in combination with (*A,B*) *mpp6*Δ, (*C,D*) *rrp47*Δ *or* (*E,F*) *rrp6*Δ. For each experiment, the wildtype *RRP40* control reveals any growth phenotype of loss of the RNA exosome cofactor. Control Wildtype cells are also shown in *A*, *C*, and *E*. *A*,*B,* While both the *rrp40-W195R mpp6*Δ and the *rrp40-Y64N mpp6*Δ double mutant cells show more impaired growth than either single mutant, the data suggest that the genetic interaction of *rrp40-Y64N* with *mpp6*Δ is stronger than the genetic interaction of *rrp40-W195R* with *mpp6*Δ as the *rrp40-Y64N mpp6*Δ cells show more severely impaired growth than the *rrp40-W195R mpp6*Δ cells (compare results in Fig. 2B to Fig. 1D). *C,D*, Both the *rrp40-W195R rrp47*Δ and the *rrp40-Y64N rrp47*Δ double mutant cells show more impaired growth than either single mutant, suggesting a genetic interaction. *E,F*, Both the *rrp40-W195R rrp6*Δ and the *rrp40-Y64N rrp6*Δ double mutant cells show more impaired growth than either single mutant, suggesting a genetic interaction. Results shown are representative of at least three independent experiments.

Analysis of *rrp40* alleles combined with loss of *RRP47* reveals that *rrp40-Y64N rrp47*Δ and *rrp40-W195R rrp47*Δ cells grow more slowly than *RRP40 rrp47*Δ cells (**Fig. 2C,D**), which are temperature-sensitive (24), or these *rrp40* single mutants at 37°C (**Fig. 1C,D**). Similarly, *rrp40-Y64N rrp6*Δ and *rrp40-W195R rrp6*Δ cells show reduced growth compared to *RRP40 rrp6*Δ cells (**Fig. 2E,F**), which are temperature-sensitive (7), or these *rrp40* single mutants at 37°C (**Fig. 1C,D**). These results indicate negative genetic interactions between *rrp40-Y64N* and *rrp40-W195R* mutants and both the *rrp47*Δ and *rrp6*Δ deletion mutants. Interestingly, *rrp40-W195R rrp47*Δ cells grow more slowly than *rrp40-Y64N rrp47*Δ cells (**Fig. 2C,D**), suggesting the *rrp47*Δ deletion exerts a greater effect on the *rrp40-W195R* mutant than the *rrp40-Y64N* mutant. None of the other *rrp40* mutants analyzed show altered growth when combined with *mpp6*Δ, *rrp47*Δ, or *rrp6*Δ. These findings suggest that among the tested *rrp40* mutants, the *rrp40-Y64N* mutant could alter RNA exosome function in a manner that that makes cells more dependent on Mpp6 function.

To analyze another critical nuclear RNA exosome cofactor, we examined genetic interactions between the series of *rrp40* mutants and variants of the essential RNA helicase Mtr4 that disrupt specific interactions required for proper Mtr4 function (71,72). For this analysis, we combined each *rrp40* mutant with the following mtr4 alleles: *mtr4-F7A-F10A*, which impairs Mtr4 interaction with Rrp6/Rrp47 (73); *mtr4-R349E-N352E*, which disrupts Mtr4 interaction with Trf4 and Air2 of the TRAMP complex while largely preserving helicase activity (74); and *mtr4-R1030A*, which lies within the helical bundle and affects Mtr4-mediated nucleic acid unwinding (75) and compared growth to cells that contain wildtype *MTR4* (**Fig. 3**). This analysis reveals that the majority of the *rrp40* mutants analyzed show no genetic interactions with any of the *mtr4* mutants tested. However, both *rrp40-W195R mtr4-R349E-N352E* and *rrp40-Y64N mtr4-R349E-N352E* double mutant cells grow more slowly than *RRP40 mtr4-R349E-N352E*, *rrp40-W195R*, and *rrp40-Y64N* single mutant cells (**Fig. 3**), revealing a specific negative genetic interaction. In contrast, both *rrp40 mtr4-F7A-F10A* and *rrp40 mtr4R-1030A* double mutant cells show no change in growth compared to the single mutant cells. This specific genetic interaction with an allele of *MTR4* that impairs interaction with the TRAMP complex provides further evidence that Rrp40 Y64N and W195R variants may impact interactions with Mpp6 as Mpp6 stimulates RNA exosome activity downstream of Mtr4/TRAMP (23,76). These findings further suggest that specific Rrp40 variants selectively impair cofactor-dependent steps in RNA surveillance rather than globally disrupting RNA exosome activity.

**Figure 3.**
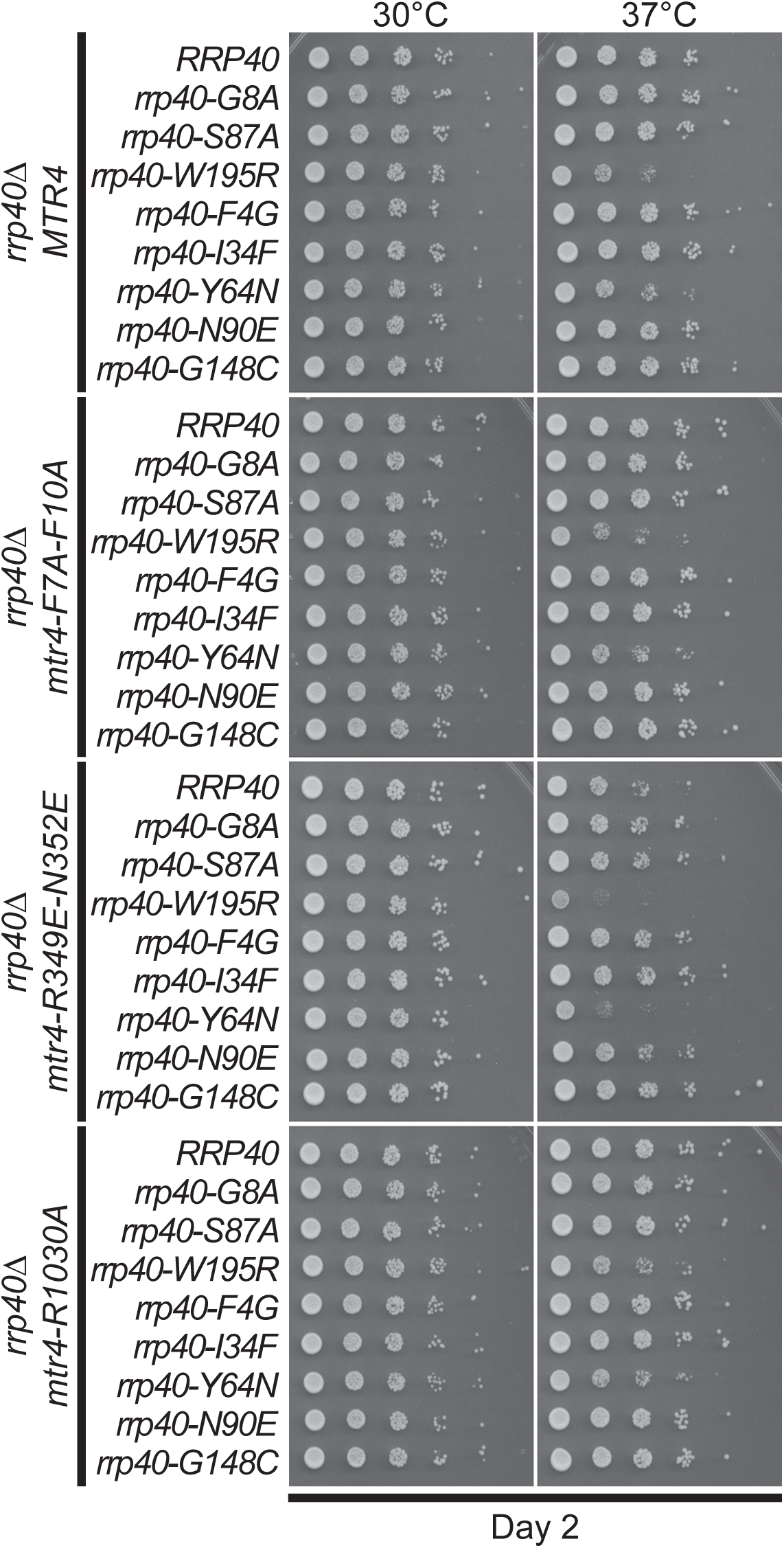
***rrp40-W195R* and *rrp40-Y64N* show genetic interactions with specific alleles of the Mtr4 RNA exosome cofactor.** Each of the *rrp40* models was combined with either control wildtype *MTR4* (*A*) or selected *MTR4* alleles that impact specific functions of Mtr4: *B*, *mtr4-F7A-F10A*, which impairs Mtr4 interaction with Rrp6/Rrp47 (73); *C*, *mtr4-R349E-N352E*, which disrupts TRAMP-mediated exosome recruitment and RNA surveillance while largely preserving helicase activity (74); or *D*, *mtr4-R1030A*, within the helical bundle, which affects Mtr4-mediated nucleic acid unwinding (75). Growth of each double mutant and an *RRP40* control that reveals any growth defect for the *mtr4* allele was analyzed by serial dilution growth assay. Results shown are representative of at least three independent experiments.

### Rrp40 variants differentially affect protein levels

While structures of the RNA exosome complex that include the cofactor Mpp6 have been reported (10,76,77), only a small region of Mpp6 is shown in these structures because most of Mpp6 is predicted to be intrinsically disordered and likely remains flexible, with only the RNA exosome-binding segment adopting a stable conformation. To more closely examine the potential consequences of how the pathogenic amino acid changes modeled could impact both the Rrp40 protein and the RNA exosome complex, we created a model of the budding yeast RNA exosome complex and Mpp6 (**Fig. 4**). We assembled a structural model of the budding yeast RNA exosome complex by integrating available cryo-EM structures with AlphaFold predictions. For the yeast RNA exosome complex, we combined the RNA exosome core (PDB ID: 6FSZ) (78) and the Mpp6-RNA exosome-RNA (PDB ID: 5VZJ) (76) structures, replacing Dis3 and reconstructing unresolved regions, including Rrp6 segments using AlphaFold2 (79). We also generated a continuous RNA substrate model by merging fragments.

**Figure 4.**
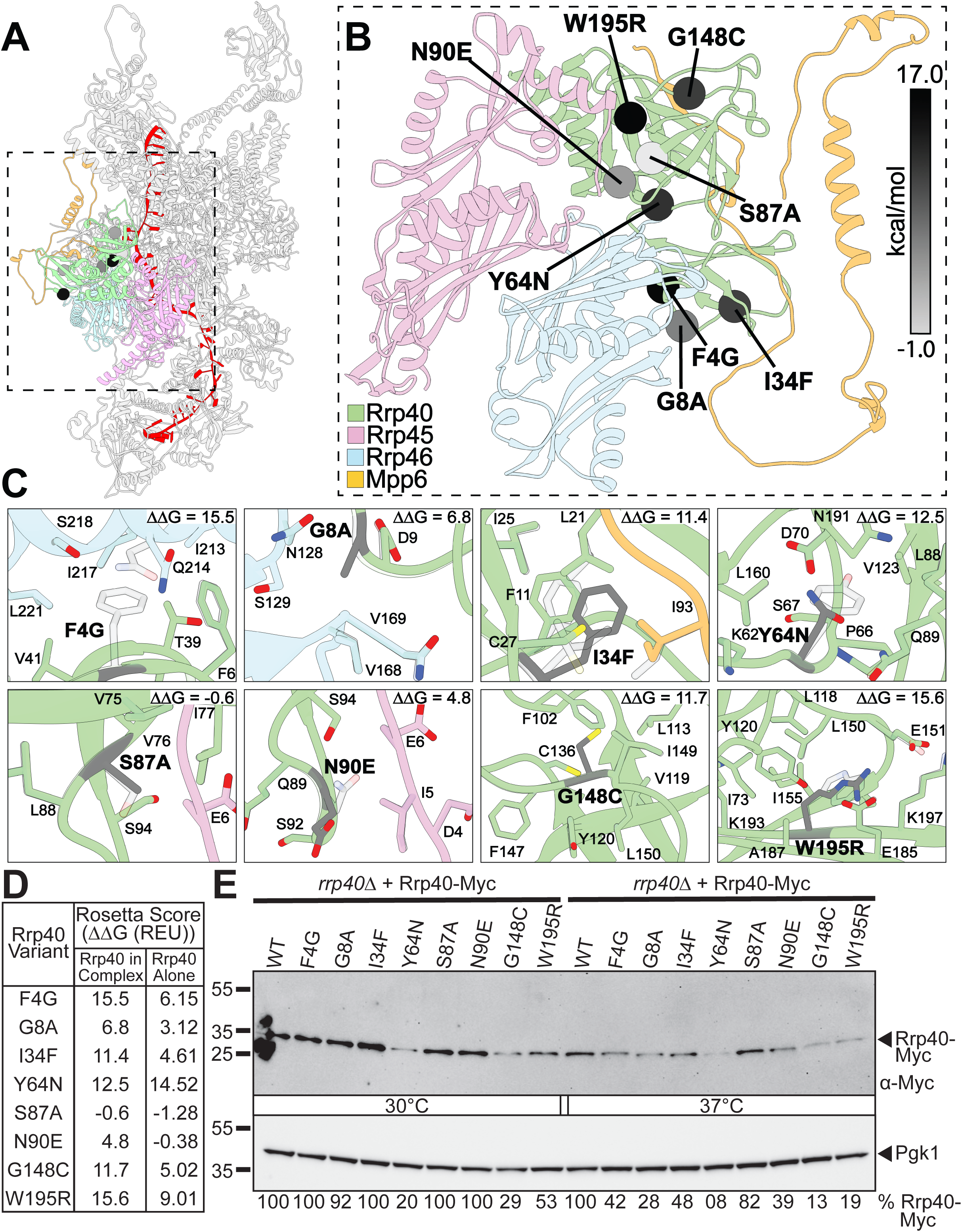
**Pathogenic Rrp40 variants are predicted to destabilize Rrp40 and the assembled RNA exosome complex**. *A*, Computational model of the budding yeast RNA exosome complex bound to RNA (red). *B*, Close-up view of the Rrp40 cap subunit (greed) and neighboring subunits Rrp45 (pink), Rrp46 (cyan), and the RNA exosome cofactor Mpp6 (orange), showing the positions of the eight residues altered in the modeled pathogenic Rrp40 variants (spheres). Spheres are shaded from white to black according to Rosetta ΔΔG value (kcal/mol). *C*, Close-up views of the local structural environment surrounding each substituted residue (grey) within the assembled complex, showing neighboring side chains and the corresponding Rosetta ΔΔG value (kcal/mol). *D*, Rosetta scores (Rosetta Energy Units-REU) calculated for each Rrp40 amino acid substitution modeled within the assembled RNA exosome complex (Rrp40 in Complex) or in the isolated Rrp40 protein (Rrp40 Alone). *E*, The steady-state level of some Rrp40 protein variants is decreased at 30°C and/or 37°C. Lysates of *rrp40*Δ cells solely expressing Myc-tagged, wildtype Rrp40 (WT) or rrp40 variants grown at 30°C or 37°C were analyzed by immunoblotting with an anti-Myc antibody to detect Rrp40-Myc and an anti-Pgk1 antibody to detect 3-phosphoglycerate kinase (Pgk1) as a loading control. Quantitation of the percentage of the rrp40 protein level relative to Rrp40 is indicated below with WT Rrp40 at each temperature set to 100%. Results are representative of two independent experiments.

As shown in **Figure 4A,B**, the Rrp40 amino acid residues that are altered in models of the pathogenic variants in EXOSC3 are located throughout the Rrp40 structure. As previously reported, Rrp40 interacts directly with Mpp6 (76,77), however, the observed genetic interaction between *rrp40-Y64N* and the *MPP6* deletion mutant (**Fig. 2A,B**) is not readily explained based on the location of Rrp40 Y64 residue relative to the position of Mpp6. To begin to assess potential structural effects of Rrp40 variants in budding yeast, Rosetta ΔΔG values were calculated for each amino acid substitution in Rrp40 both in the context of the isolated protein and within the RNA exosome complex (**Fig. 4C,D**). Overall, most Rrp40 substitutions are predicted to be destabilizing, with generally enhanced effects when Rrp40 is evaluated in the context of the assembled RNA exosome complex (**Fig. 4C,D**), indicating that the packing constraints and intersubunit contacts within the complex leave less tolerance for local structural perturbations than the isolated protein alone. Several Rrp40 variants predict strong destabilizing effects when modeled within the complex, including F4G (ΔΔG: 15.5 kcal/mol), W195R (15.6), Y64N (12.5), G148C (11.7), and I34F (11.4). These variants show consistently positive ΔΔG values, indicating substantial destabilization of Rrp40 in the assembled context. For the isolated Rrp40 protein, the predicted destabilizing effects of several of these variants were more modest, as seen for F4G (6.1), G148C (5.0), and I34F (4.6), suggesting that the local environment of the complex, including packing against neighboring subunits, amplifies the structural cost of these substitutions rather than absorbing this destabilization. For instance, F4G replaces a bulky aromatic phenylalanine side chain with glycine, which lacks a side chain; this loss of steric bulk and hydrophobic packing may be reasonably well tolerated when Rrp40 is isolated and conformationally flexible but may become far more disruptive once the residue is constrained at a fixed interface within the complex, where the resulting cavity cannot be compensated for.

The Rrp40-Y64N variant is predicted to be strongly destabilizing both within the context of the isolated protein and within the complex, with a ΔΔG of 14.5 calculated for the isolated protein and ΔΔG of 12.5 for the complex-embedded protein, indicating that the contribution of this residue to local stability is largely intrinsic rather than dependent on the surrounding complex architecture. This prediction of the model is consistent with the central location of Y64 residue relative to other RNA exosome ring subunits, Rrp45 and Rrp46 (**Fig. 4B**). The tyrosine-to-asparagine substitution changes a large aromatic side chain capable of hydrophobic and potentially π-stacking contacts to a much smaller, polar amide-containing side chain. This change is likely to disrupt local packing or hydrogen-bonding networks regardless of whether Rrp40 is examined alone or in the context of adjacent subunits.

Rrp40 variants with moderate effects in the complex predict reduced destabilization or modest behavior compared to these variants modeled in isolation, such as G8A (ΔΔG: 3.1 isolated; 6.8 in complex) and N90E (ΔΔG: 0.4 isolated; 4.8 in complex; −0.4 alone), further supporting a model in which the environment within the complex rather than the isolated protein, is more sensitive to these substitutions. Notably, two substitutions, S87A (ΔΔG: −1.3 alone; −0.6 in complex) and N90E (-0.4 alone), are predicted to have neutral or slightly stabilizing effects in isolation, suggesting minimal intrinsic structural disruption outside the context of the complex. Collectively, these results indicate that while several of the Rrp40 variants are predicted to be only mildly destabilizing on their own, incorporation into the RNA exosome complex likely often exacerbates their energetic impact, consistent with these residues making contacts or occupying packing environments that are only present once Rrp40 is assembled with its partner subunits.

To experimentally test the predictions from the Rosetta protein stability analysis, we performed immunoblotting on C-terminally Myc-tagged wildtype Rrp40 and Rrp40 variants expressed as the sole copy of Rrp40 in yeast cells grown at 30°C and 37°C (**Fig. 4E,S1**). In cells grown at 30°C, three Rrp40 variants show a decrease in steady-state protein levels compared to wildtype Rrp40: Rrp40 Y64N (20% relative to WT); Rrp40 G148C (29% relative to WT); and Rrp40 W195R (53% relative to WT) (**Fig. 4E**). In cells grown at 37°C, most of the Rrp40 variants analyzed show some decrease in steady-state protein levels relative to wildtype Rrp40, particularly Rrp40 Y64N (8% relative to WT), Rrp40-G148C (13% relative to WT), and Rrp40-W195R (19% relative to WT) (**Fig. 4E**). The observed decreases in the steady-state levels of the Rrp40 variants at 37°C correlate quite well with the predicted decreases in the stabilities (positive Rosetta scores) of the variants within the complex (**Fig. 4D**).

Many of the Rrp40 variants that show a decrease in steady-state levels, including Rrp40 G148C, which shows a decrease similar to Rrp40 W195R (**Fig. 4E**), cause no detectable growth defect (**Fig. 1C,D**) or show no negative genetic interactions (**Figs. 2,3**) at 37°C. This finding indicates that reduced steady-state protein levels of Rrp40 variants do not consistently correlate with cellular phenotypes, suggesting that decrease in protein levels alone is insufficient to explain the observed growth and genetic interaction defects. Instead, these results support a model in which specific Rrp40 variants likely impair discrete structural or functional features of the RNA exosome complex, beyond global protein stability, that underlie functional defects.

### Rrp40 Y64N impairs RNA exosome function

Given the finding that the Rrp40 Y64N variant, modeling EXOSC3, which has not previously been analyzed, shows temperature-sensitive growth defects and negative genetic interactions with both *MPP6* and *MTR4*, we used the CRISPR/Cas system to edit the endogenous *RRP40* locus and generate *rrp40-Y64N* cells (**Fig. 5**). Three independent *rrp40-Y64N* isolates show similar growth defects (**Fig. 5A**). While *rrp40-Y64* cells show slow growth compared to control wildtype cells, *rrp40-W195R* cells generated by the same CRISPR/Cas genome editing strategy (50), show slower growth than *rrp40-Y64N* (**Fig. 5A**). The slow growth phenotype of the three independent *rrp40-Y64N* isolates is rescued by a plasmid-borne copy of wildtype *RRP40* (**Fig. 5B**), confirming that the slow growth phenotype of the newly generated *rrp40-Y64N* model is due to the specific engineered mutation in *RRP40*.

**Figure 5.**
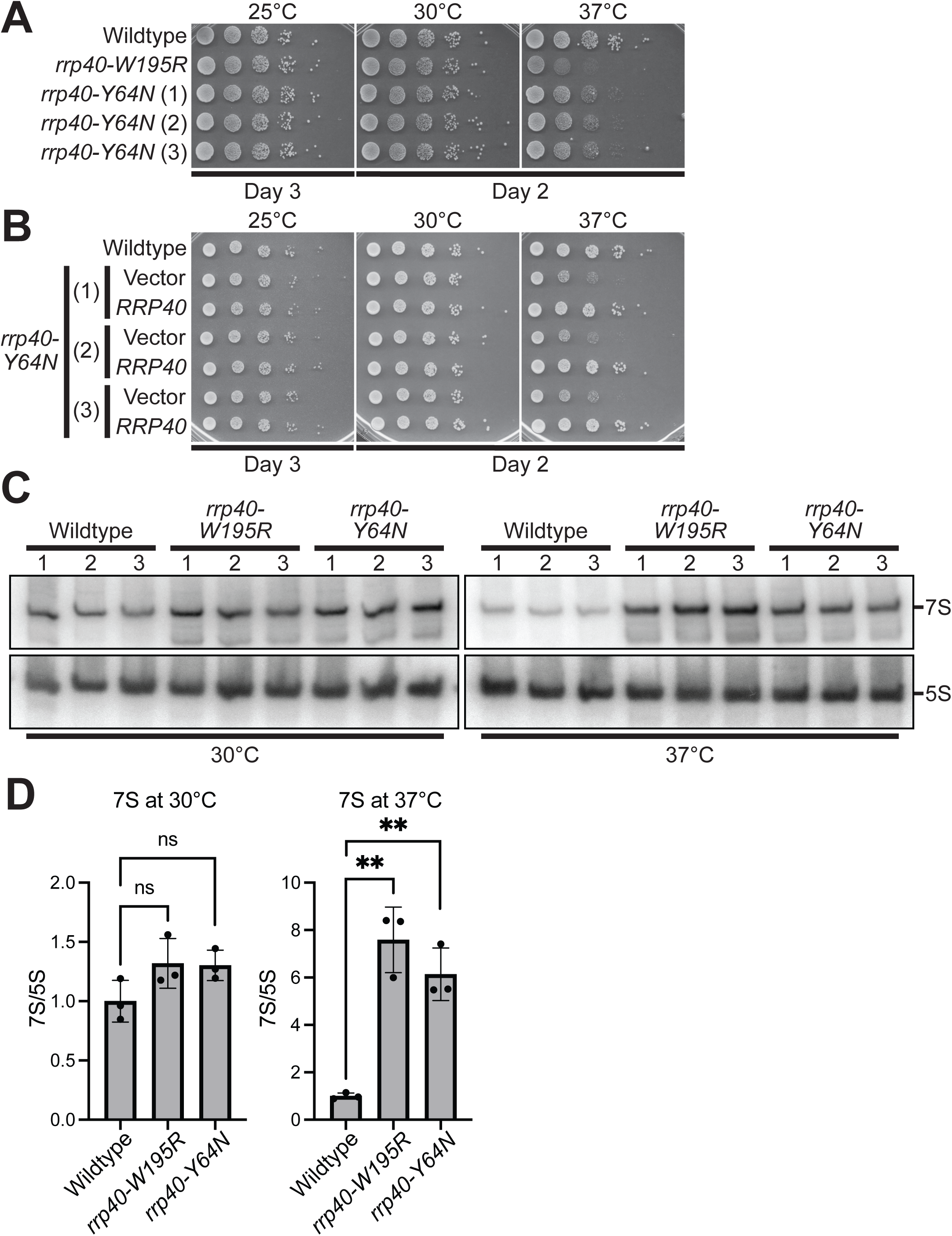
**The *rrp40-Y64N* mutant cells exhibit defects in rRNA processing**. *A*, CRISPR/Cas-generated models of rrp40-W195R, which serves as a control because this model has been analyzed previously (52), and rrp40-Y64N show growth defects at 37°C as assessed by a serial dilution growth assay. Three independent CRISPR/Cas-generated mutants (1–3) are shown for *rrp40-Y64N*. *B*, The three *rrp40-Y64N* CRISPR mutant isolates containing Vector or *RRP40* plasmid and a Wildtype control were serially diluted, spotted, and grown at the indicated temperatures for 2-3 days, demonstrating that the temperature-sensitive growth is due to the *rrp40* allele. *C*,*D*, Northern blotting reveals rRNA processing defects evident as accumulation of 7S pre-rRNA in *rrp40-Y64N* mutant cells. Equal amounts of total RNA from *RRP40, rrp40-W195R*, and *rrp40-Y64N* mutant cells grown at either 30°C or 37°C were used to analyze levels of 7S pre-rRNA by northern blotting. The 5S rRNA was probed as a loading control. Experiments were performed in triplicate (labeled 1,2, and 3). *D*, The blots shown in (*C*) were quantitated by analyzing the ratio of 7S pre-rRNA to 5S RNA and setting this ratio to 1.0 in *RRP40* control cells at each temperature. Statistical significance was calculated by t-test (**P-value ≤ 0.01; ns not significant).

To assess whether the Rrp40 Y64N variant impairs the function of the RNA exosome, we compared rRNA processing in *rrp40-Y64N* cells to *rrp40-W195R* cells, which have been previously analyzed (50,52). For this analysis, total RNA from Wildtype, *rrp40-W195R*, and *rrp40-Y64N* cultures grown at 30°C and 37°C was analyzed by northern blotting with probes to detect 7S pre-rRNA and 5S rRNA (**Fig. 5C**). The 7S pre-rRNA is a direct substrate of the nuclear RNA exosome (4,12,80). As shown in **Figure 5C** and quantitated in **Figure 5D**, at 37°C, *rrp40-Y64N* cells accumulate the 7S rRNA precursor to nearly the same extent as *rrp40-W195R* cells. We also examined a set of well-defined target non-coding RNAs (80–82) that are processed RNA exosome using the *rrp40-W195R* and *rrp40-Y64N* plasmid shuffle cells (**Fig. S2**). Total RNA from wildtype and *rrp40* cells grown at 37°C was analyzed by RT-qPCR with primers to detect *U4* pre-snRNA, *TLC* pre-RNA, and *U14* snoRNA. We observed a statistically significant increase in the steady state levels of *U4* pre-snRNA (**Fig. S2A**) and *TLC1* pre-RNA (**Fig. S2B**) for both the *rrp40-Y64N* and *rrp40-W195R* cells. A statistically significant increase in the steady-state level of *U14* snoRNA was only observed in the *rrp40-W195R* cells (**Fig. S2C**). In all cases, the level of RNA exosome target RNAs accumulated in *rrp40-W195R* cells was greater than detected in *rrp40-Y64N* cells, consistent with the more significant growth defect in *rrp40-W195R* cells. Together, these studies provide the first evidence that the Rrp40 Y64N variant modeling EXOSC3 Y109N impairs RNA exosome function.

### Rrp40 W195R and Y64N variants impair functional interaction of the RNA exosome with Mpp6

The results described here for both the *rrp40-Y64N* and *rrp40-W195R* cells and previous biochemical and structural analysis of Rrp40 W195R (76,77) suggest that these amino acid substitutions could impair interaction with the RNA exosome cofactor Mpp6. As a test of this prediction, we examined whether overexpression of wildtype *MPP6* from a high copy plasmid (2μ) suppresses the growth defect of *rrp40-W195R* and *rrp40-Y64N* cells. Results of this experiment show that overexpression of *MPP6* can suppress the temperature-sensitive growth of both *rrp40-W195R* and *rrp40-Y64N* cells relative to cells with vector alone at 37°C (**Fig. 6A**).

**Figure 6.**
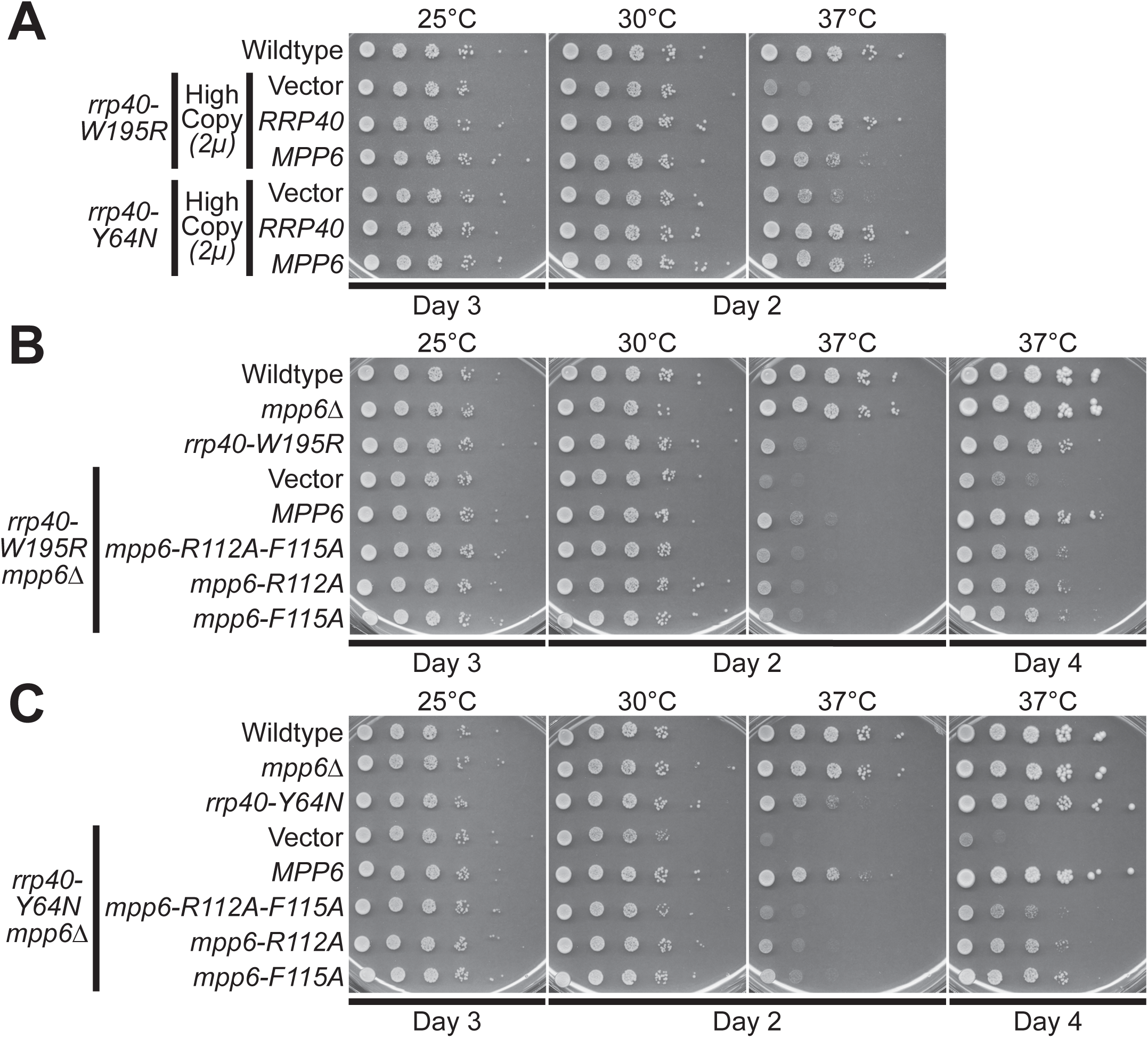
Overexpression of Mpp6 can rescue growth defects of *rrp40-W195R* and *rrp40-Y64N* mutant cells. *A*, Plasmids that overexpress either Rrp40, as a control, or Mpp6 were transformed into *rrp40-W195R* or *rrp40-Y64N* mutant cells and growth at the indicated temperatures was assayed by serial dilution growth assay. Wildtype cells are also included as a control. *B*, Mpp6 interaction with the Rrp40 is required to rescue growth defects of *rrp40* mutant cells. *B*, *rrp40-W195R* or *C*, *rrp40-Y64N* cells also lacking Mpp6 (i.e., *rrp40 mpp6*Δ double mutant cells, which lack endogenous Mpp6) were transformed with Vector or plasmids encoding wildtype *MPP6*, *mpp6-R112A-F115A*, *mpp6-R112A*, or *mpp6-F115A*. Mpp6 R112 and F115 are key residues required for functional Mpp6 interaction with Rrp40 and stimulation of RNA exosome activity (76,77). These transformed cells as well as WT control, *mpp6*Δ, and *rrp40-W195R* cells were analyzed by serial dilution growth assay for the indicated number of Days at 25°C, 30°C or 37°C. Day 2 and Day 4 growth are shown for at 37°C to better visualize suppression of temperature sensitive growth, which is lost for the Mpp6 variants that have impaired interaction with Rrp40. Results shown are representative of at least three independent experiments.

Previous studies identified Mpp6 R112 and F115 as key residues required for functional interaction of Mpp6 with Rrp40 and stimulation of RNA exosome activity (76,77). We tested whether the Mpp6 variants, Mpp6 R112A, F115A, Mpp6 R112A, or Mpp6 F115A interfere with Mpp6-mediated rescue of *rrp40-W195R* (**Fig. 6B**) or *rrp40-Y64N* (**Fig. 6C**) temperature sensitive growth when Mpp6 is expressed from a low copy plasmid (*CEN*) in cells lacking endogenous *MPP6* (*rrp40 mpp6*Δ). These studies reveal that this low level of *MPP6* expression can rescue the temperature sensitive growth defect of both *rrp40-W195R* (**Fig. 6B**) and *rrp40-Y64N* (**Fig. 6C**) as compared to Vector control. Furthermore, Mpp6 R112A, F115A impairs this Mpp6-mediated rescue for both *rrp40-W195R* (**Fig. 6B**) and *rrp40-Y64N* (**Fig. 6C**). Either single amino acid substitution alone also impairs the Mpp6-mediated rescue but not to the same extent as Mpp6 R112A, F115A, which contains both amino acid substitutions. We also further confirmed these results in *rrp40-Y64N* cells demonstrating that the overexpression (2μ plasmid) of wildtype Mpp6 but not Mpp6 R112A, F115A, Mpp6 R112A, or Mpp6 F115A can suppress temperature sensitive growth even when the cells contain an endogenous copy of *MPP6* (**Fig. S3**).

These results demonstrate that the ability of Mpp6 to suppress the growth defects of Rrp40 variants depends on residues previously shown to mediate direct interaction of this cofactor with Rrp40. The reduced rescue/suppression observed with Mpp6 interface variants, particularly the R112A, F115A double substitution, indicates that efficient functional rescue requires the Rrp40-Mpp6 interaction. These data are consistent with altered functional interaction of these Rrp40 variants with Mpp6.

### Pathogenic EXOSC3 variants differentially affect protein levels

To extend this analysis to human EXOSC3, we created a structural model of the human RNA exosome including MPP6 as described in Experimental procedures (**Fig. 7**). For the human complex, we used the nuclear RNA exosome-MTR4-RNA structure (PDB: 6D6R) (83) as a framework and positioned MTR4 and EXOSC10 using AlphaFold (79). The MTR4-EXOSC10 module was fitted into the cryo-EM structure, while remaining regions and missing loops across RNA exosome subunits were modeled with AlphaFold to produce a complete complex. Similar to budding yeast Rrp40 (**Fig. 4B**), the amino acids altered in disease are spread across the EXOSC3 protein rather than being localized at a specific hot spot (**Fig. 7B**).

**Figure 7.**
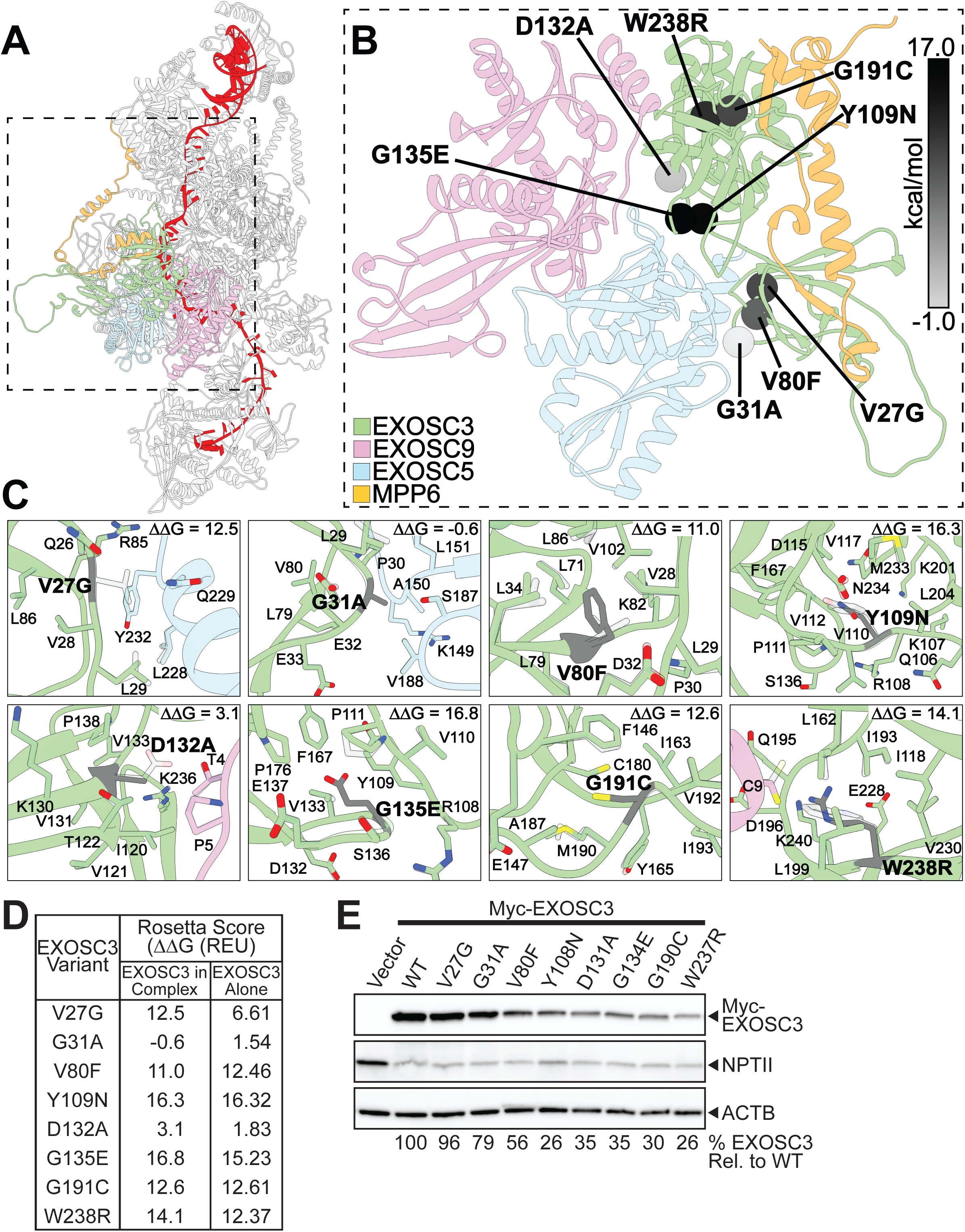
**Pathogenic EXOSC3 variants are predicted to destabilize EXOSC3 and the assembled RNA exosome complex**. *A*, Computational model of human RNA exosome complex bound to RNA (red). *B*, Close-up view of the EXOSC3 cap subunit (green) and neighboring subunits EXOSC9 (pink), EXOSC5 (cyan), and the RNA exosome cofactor MPP6 (orange), showing the positions of the eight residues altered in the pathogenic EXOSC3 variants (spheres). Spheres are shaded from white to black according to Rosetta ΔΔG value (kcal/mol). *C*, Close-up views of the local structural environment surrounding each substituted residue (grey), within the assembled complex, showing neighboring side chains and the corresponding Rosetta ΔΔG value (kcal/mol) calculated for that substitution in complex. *D*, Rosetta scores (Rosetta Energy Units-REU) calculated for each EXOSC3 amino acid substitution modeled within the assembled RNA exosome complex (ECOSC3 in Complex) or in the isolated ECOSC3 protein (EXOSC3 Alone). *E*, The steady-state level of some of the EXOSC3 protein variants is decreased compared to wildtype (WT) EXOSC3. Lysates of mouse N2a cells transfected with empty vector, vector expressing murine Myc-EXOSC3 or the indicated Myc-EXOSC3 variants (murine residue numbering, one residue lower than the corresponding human EXOSC3 position) were analyzed by immunoblotting with anti-Myc antibody to detect Myc-EXOSC3 proteins. ACTB serves as a loading control and neomycin phosphotransferase II (NPTII) serves as a transfection control. Quantitation of the percentage of the EXOSC3 protein level relative to EXOSC3 variants is indicated below, with WT Myc-EXOSC3 set to 100%. Results are representative of two independent experiments.

Using the model generated based on the available structural information, we evaluated the energetic effects of EXOSC3 substitutions by calculating Rosetta ΔΔG values for each variant in both the isolated EXOSC3 protein and within the assembled RNA exosome complex (**Fig. 7C**). Most variants are predicted to be destabilizing, with several showing consistent effects in both isolated and complex contexts (**Fig. 7D**). Strong destabilizing substitutions include Y109N (ΔΔG: 16.3 alone; 16.3 in complex), G135E (16.8 alone; 15.2 in complex), W238R (14.1 alone; 12.4 in complex), G191C (12.6 alone; 12.6 in complex), and V80F (11 alone; 12.5 in complex), indicating that these effects are largely intrinsic to EXOSC3 and are not substantially buffered by incorporation into the RNA exosome complex. In contrast, variants such as V27G and D132A show reduced destabilization in the complex, suggesting partial stabilization through protein-protein interactions, while G31A displays minimal and context-dependent effects. The V27G variant (6.6 alone; 12.5 in complex) shows increased destabilization in the complex, suggesting partial destabilization through protein-protein interactions. Comparison of the yeast and human models and the variant stabilities for Rrp40 vs EXOSC3 (**Figs. 4, 7**) indicates that while incorporation into the RNA exosome complex generally increases the destabilizing effects of Rrp40 variants in yeast, many human EXOSC3 variants show similar strong destabilizing effects in isolation and in complex, suggesting overall variant intrinsic misfolding or instability.

To experimentally test the EXOSC3 variant stability predictions, we performed immunoblotting on N-terminally Myc-tagged wildtype mouse EXOSC3 and EXOSC3 variants expressed in a mouse neuronal cell line, Neuro-2a (N2a) cells (84) (**Fig. 7E**). Many of the EXOSC3 variants show a decrease in steady-state protein levels, supporting most of the model stability predictions. In particular, mouse EXOSC3 Y108N and W237R, corresponding to human EXOSC3 Y109N and W238R variants, show the most pronounced reduction (26% relative to WT). Interestingly, the extent of reduction in protein levels correlates with variant position, with substitutions closer to the C-terminus showing greater decreases in protein abundance. Together, these results suggest that disease-associated EXOSC3 variants, including the previously uncharacterized Y109N variant, alter protein stability in mammalian cells.

### EXOSC3 Y109N disrupts RNA exosome complex interactions

To experimentally test whether EXOSC3 variants disrupt assembly/integrity of the mammalian RNA exosome complex, we examined interactions between EXOSC3 variants and RNA exosome subunits.

Wither wildtype (WT) Myc-tagged mouse EXOSC3 or the indicated EXOSC3 variants were expressed in mouse N2a cells and subjected to immunoprecipitation, followed by immunoblotting to detect associated endogenous RNA exosome subunits, including the core ring subunits EXOSC8 and EXOSC9, as well as the catalytic subunit EXOSC10 (**Fig. 8**). Histone H3 was probed as a loading control. This analysis shows that mouse EXOSC3 Y108N variant, corresponding to human EXOSC3 Y109N, exhibits a marked reduction in interaction with all tested subunits, suggesting a substantial defect in incorporation into our stability within the RNA exosome complex. The loss of interaction was observed across both core ring and associated EXOSC10 subunits, suggesting that this Y109N substitution broadly disrupts RNA exosome assembly/integrity rather than affecting a single interface. In contrast, the other EXOSC3 variants analyzed show interaction profiles similar to wildtype EXOSC3, despite several exhibiting reduced steady-state protein levels (**Fig. 7**).

**Figure 8.**
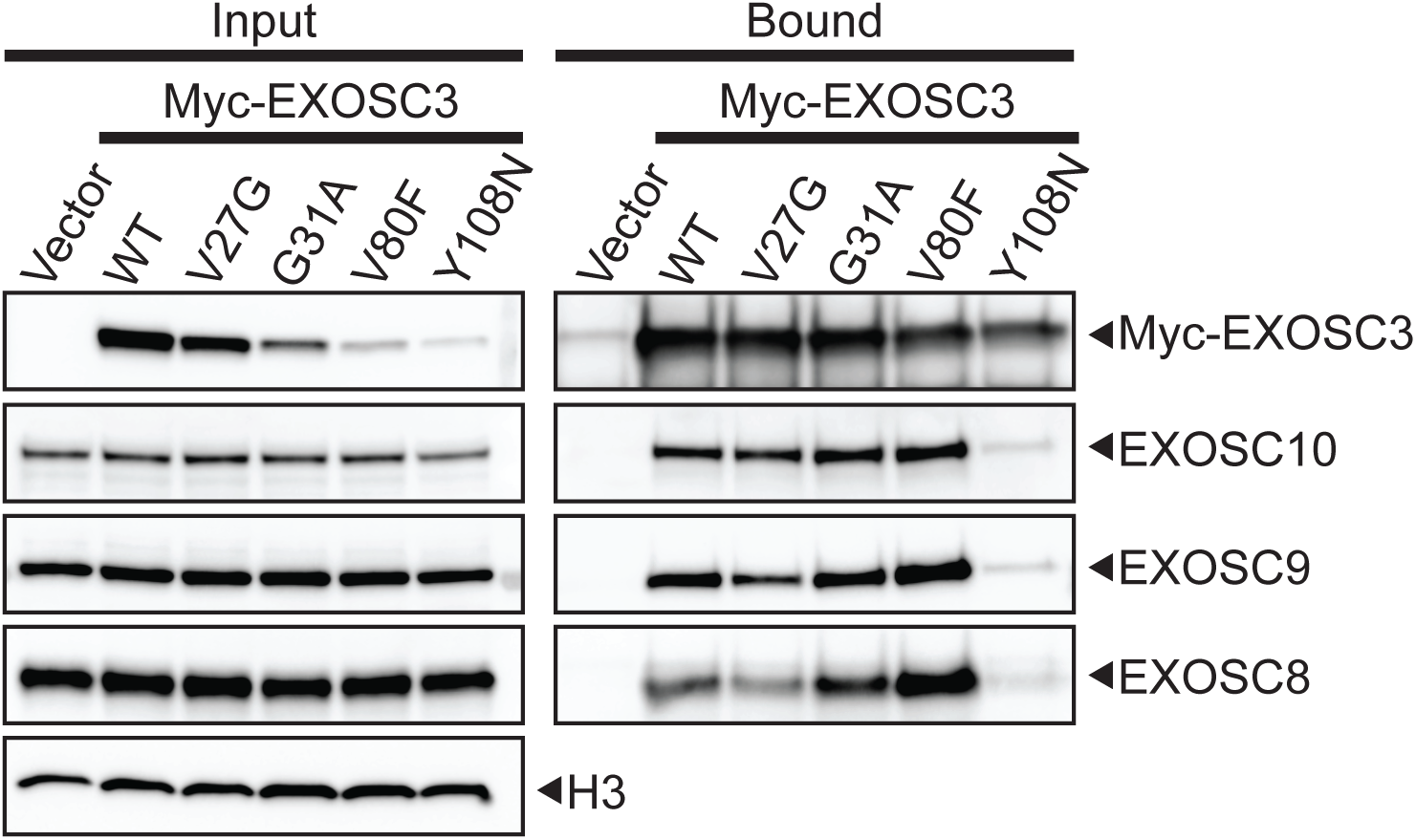
The pathogenic amino acid substitutions in EXOSC3 can alter interactions with other RNA exosome subunits. *A*, Myc-EXOSC3 or Myc-EXOSC3 variants were immunoprecipitated from N2A cells and interactions with the RNA exosome subunits EXOSC8, EXOSC9, and the RNA exosome-associated subunit EXOSC10 were analyzed by immunoblotting. Both the Input and Bound samples are shown. NPTII represents the neomycin phosphotransferase II, which is encoded on the Myc-EXOSC3 and Myc-EXCOSC3 variant plasmids, indicating similar levels of transfection across the EXOSC3 variants. H3 serves as a loading control for the Input sample. Results are representative of at least three independent experiments.

These findings indicate that for the mouse EXOSC3 G31A and V80F variants, decreased protein abundance does not necessarily impair their ability to associate with the RNA exosome. Instead, the G31A and V80F variants likely impact protein stability without strongly affecting complex assembly/integrity. These results also suggest that EXOSC3 variants differentially affect RNA exosome assembly: the mouse EXOSC3 Y108N variant modeling human EXOSC3 Y109N disrupts interactions with multiple RNA exosome subunits, in contrast to mouse G31A and V80F EXOSC3 variants that primarily reduce protein stability while preserving subunit interactions.

## Discussion

In this study, we performed a systematic analysis of disease-associated EXOSC3 missense variants using complementary budding yeast and mammalian model systems. By integrating genetic, biochemical, and computational approaches, we identify variant-specific mechanisms by which EXOSC3 variants impact RNA exosome function. Collectively, our findings demonstrate that pathogenic EXOSC3 variants do not impair function through a single uniform mechanism but instead impair RNA exosome activity through distinct effects on protein stability, cofactor interactions, and complex assembly/stability.

A key conclusion of our work is that many EXOSC3 variants are likely intrinsically destabilizing but decrease in protein levels alone is not sufficient to explain the observed functional or cellular phenotypes of these variants. Both computational stability predictions and immunoblot analyses indicate that several variants reduce steady-state protein levels in budding yeast and mammalian cells. However, several of these variants that show substantial decreases in steady-state protein level do not show measurable growth defects or strong genetic interactions in yeast. This disconnect indicates that moderate reductions in EXOSC3 stability, and by extension, total available RNA exosome levels, can be tolerated without severely compromising essential RNA processing functions. These findings are consistent with prior studies (41,49), suggesting that the RNA exosome retains functional activity, even when subunit levels are partially reduced. Overall, while decreased subunit stability likely contributes to disease pathology, this change cannot fully account for the phenotypic diversity observed in patients.

Our data support a model in which specific amino acid substitutions disrupt distinct functional features of the RNA exosome. This is most clearly demonstrated by the case of Y109N variant. In both yeast and mammalian systems, this variant shows a combination of defects that distinguish it from other variants. In yeast, *rrp40-Y64N* causes temperature-sensitive growth, and measurable defects in rRNA and other ncRNA processing and shows genetic interactions with both *MPP6* and *MTR4*. In mammalian cells, the corresponding EXOSC3 Y109N variant shows reduced protein levels and, importantly, a clear defect in interaction with other RNA exosome subunits. This loss of interaction suggests a failure to properly assemble into or maintain the integrity of the RNA exosome complex. Together, these findings identify Y109N as a mechanistically distinct variant that likely impairs both protein stability and complex assembly/integrity, providing a clear example of how individual pathogenic amino acid variants can impair RNA exosome function through multiple converging defects.

Our analysis also provides new insight into the role of RNA exosome cofactors in modulating the functional impact of EXOSC3 variants. Genetic interaction studies in yeast reveal that *rrp40-Y64N* and *rrp40-W195R* show specific sensitivity to loss of Mpp6 function and to mutations in *MTR4* that disrupt TRAMP-mediated recruitment. These interactions point to a functional defect in cofactor-dependent RNA processing pathways rather than a global loss of RNA exosome activity. Consistent with this interpretation, overexpression of Mpp6 suppresses the growth defects of both *rrp40-Y64N* and *rrp40-W195R*, and this suppression requires residues in Mpp6 that mediate direct interaction with Rrp40. These findings strongly support a model in which certain disease-associated variants weaken or disrupt productive engagement between the RNA exosome and key cofactors. Given that cofactors such as Mpp6 and Mtr4 play essential roles in substrate targeting and activation of RNA exosome activity (22,69,76,77,83), even subtle changes in these interactions could have significant downstream effects on RNA metabolism.

Comparison of the budding yeast and human structural analyses further highlights important differences in how the RNA exosome complex responds to destabilizing amino acid substitutions. In the yeast system, incorporation of Rrp40 into the RNA exosome complex frequently sensitizes the energetic impact of destabilizing variants, consistent with packing constraints and intersubunit contacts within the complex allowing less tolerance for local structural perturbations. In contrast, several human EXOSC3 variants show strong destabilizing effects for the subunit in isolation that are retained even in the context of the full RNA exosome complex. The reduced buffering capacity in the yeast system may reflect differences in the structural organization or interaction networks of the complex, supporting the possibility that certain human EXOSC3 variants may impose persistent folding/stability defects that cannot be compensated for by assembly into the complex. Notably, the EXOSC3 variants that produce the most pronounced defects when modeled in budding yeast, EXOSC3 Y109N and W238R, have not been reported as homozygous in any clinical report to date (46), consistent with the hypothesis that major loss of RNA exosome function is incompatible with life. These observations support a model in which pathogenic mechanisms can arise from both context-dependent disruption of RNA exosome architecture and intrinsic defects in subunit folding, highlighting the complementary insights gained from yeast and mammalian systems

The diversity of molecular defects observed, ranging from decreased protein levels to impaired cofactor interactions and defective complex assembly/integrity, suggests that different variants may selectively compromise distinct RNA processing pathways. Neurons, and particularly the specific cell types that comprise the cerebellum, may be particularly sensitive to such perturbations due to their high dependence on precise RNA processing and surveillance mechanisms. Thus, even modest or pathway-specific disruptions of RNA exosome function could lead to the selective vulnerability observed in pontocerebellar hypoplasia and related disorders. Recent studies employing cerebellar organoids have begun to provide evidence in support of a model where specific cells types are more dependent on the function of the RNA exosome than other cells types (56). From a broader perspective, this work emphasizes the importance of allele-specific functional characterization in understanding the molecular basis of genetic disease. Our results highlight that even closely located amino acid substitutions within the same structural subunit of a macromolecular complex, such as EXOSC3 V27G and G31A, can have markedly different biochemical and functional consequences. This systematic analysis reinforces the need to move beyond simple genotype-phenotype correlations and toward detailed mechanistic analyses of individual variants.

The observation that Mpp6 overexpression can suppress growth defects associated with specific Rrp40 variants in budding yeast suggests that defining the molecular consequences of individual disease-associated variants can provide insight into potential mechanisms that modulate RNA exosome function. These findings raise the possibility that alterations in cofactor interactions or pathways that enhance RNA exosome activity may influence the functional impact of certain variants. Additionally, variants that primarily affect protein stability or complex assembly may be differentially responsive to conditions that block protein degradation or promote protein stability. While these observations are derived from a yeast model system and will require further validation in more disease-relevant contexts, the findings highlight the importance of variant-specific mechanisms in shaping RNA exosome dysfunction and may help guide future studies aimed at understanding potential avenues for modulating RNA exosome activity in disease.

In summary, this systematic analysis reveals that disease-associated EXOSC3 variants disrupt RNA exosome function through distinct mechanisms. By combining insights from genetics, functional assays, and structural modeling across two model systems, we provide a framework for understanding how specific amino acid substitutions within one subunit of the essential RNA exosome complex may contribute to disease etiology.

## Experimental procedures

### Chemicals and media

All chemicals were obtained from Sigma-Aldrich, United States Biological, or Fisher Scientific unless otherwise noted. All media were prepared by standard procedures (85).

### Protein structure analysis

We used the structures of human [PDB: 6D6R; (83)] and budding yeast RNA exosome complex [PDB: 6FSZ; (78)]. The PyMOL Molecular Graphics System, Version 2.0 Schrödinger, LLC was used for viewing structures and figure making. AlphaFold was used for structure predictions of the variants (79).

### S. cerevisiae strains and plasmids

All DNA manipulations were performed according to standard procedures (86). *S. cerevisiae* strains and plasmids used in this study are listed in **Table S2** and oligonucleotides are listed in **Table S3**. The *rrp40*Δ (yAV1107) haploid yeast strain was generated as previously described (87). The *rrp40*Δ *mpp6*Δ (ACY2638), *rrp40*Δ *rrp47*Δ (ACY2462), *rrp40*Δ *rrp6*Δ (ACY2466), and *rrp40*Δ *mtr4*Δ (ACY2540) haploid yeast strains were generated as previously described (88). The *rrp40-W195R* (ACY3117) CRISPR mutant strain was generated as previously described (50). Two additional *rrp40-W195R* (ACY3118-3119) CRISPR mutant strains employed in this work were generated identically to ACY3117 (50). The *mpp6*Δ (ACY3315), *rrp40-W195R mpp6*Δ (ACY3321), and *rrp40-Y64N mpp6*Δ (ACY3327) haploid yeast strains were constructed by deletion of the *MPP6* ORF in the wildtype (BY4741), *rrp40-W195R* (ACY3117), and *rrp40-Y64N* (ACY3290) strains, respectively, by homologous recombination using *MPP6-UTR-natMX4* PCR products amplified from *natMX4* cassette plasmid [p4339; (89)] and oligonucleotides (AC7967/7968; Integrated DNA Technologies).

The wildtype *RRP40* (pAC3652), *rrp40-S87A* (pAC3654), *rrp40-W195R* (pAC3655) *CEN6 LEU2* plasmids, which contain the endogenous *RRP40* promoter, 5’UTR, and 3’UTR, were constructed as previously described (50,88). The *RRP40-2xMyc* (pAC3666) *CEN6 LEU2* plasmid was generated by excision of the *2×Myc-ADH1* 3′-UTR from the *RRP40-2xMyc-ADH1* 3’-UTR (pAC3161) *CEN6 LEU2* plasmid (51) by restriction digestion and cloning of the native *2×Myc-RRP40* 3′-UTR PCR product into the plasmid using NEBuilder HiFi Assembly (New England BioLabs). The *rrp40-F4G* (pAC4515), *rrp40-G8A* (pAC3653), *rrp40-I34F* (pAC4516), *rrp40-Y64N* (pAC4517), *rrp40-N90E* (pAC4518), and *rrp40-G148C* (pAC4519) *CEN6 LEU2* plasmids were generated by site-directed mutagenesis of the *RRP40* (pAC3652) *CEN6 LEU2* plasmid using oligonucleotides encoding each amino acid change (AC5187-5188; AC10172-10179; Integrated DNA Technologies; **Table S3**) and QuickChange II SDM Kit (Agilent). The *rrp40-F4G-2xMyc* (pAC4521), *rrp40-G8A-2xMyc* (pAC4520), *rrp40-I34F-2xMyc* (pAC4522), *rrp40-Y64N-2xMyc* (pAC4523), *rrp40-S87A-2xMyc* (pAC3667), *rrp40-N90E-2xMyc* (pAC4524), *rrp40-G148C-2xMyc* (pAC4525), and *rrp40-W195R-2xMyc CEN6 LEU2* plasmids were generated by site-directed mutagenesis of the *RRP40-2xMyc* (pAC3666) *CEN6 LEU2* plasmid using oligonucleotides encoding each amino acid change (AC5187-5188; AC5982-5985; AC10172-10179; Integrated DNA Technologies; **Table S3**) and QuickChange II SDM Kit (Agilent). The *MTR4* (pAC4096), *mtr4-F7A-F10A* (pAC4099), *mtr4-R349E-N352E* (pAC4100), and *mtr4-R1030A* (pAC4104) *CEN6 HIS3* plasmids were generated as previously described (88,90). The *TEF1p-Cas9-CYC1t-SNR52p* (pAC3846) *CEN6 URA3* pCas9 plasmid was constructed as previously described (50). The *TEF1p-Cas9-CYC1t-SNR52p-RRP40_164.gRNA-SUP4t* (pAC4560) *CEN6 URA3* pCas9 plasmid containing the gRNA for targeting *RRP40* was constructed by PCR amplification of *RRP40_164.gRNA* with oligonucleotides AC10660 and AC10140 (Integrated DNA Technologies) using p426-*SNR52p-gRNA.CAN1.Y-SUP4t* plasmid template [Addgene #43803, (91)] and cloning of gRNA PCR product into pAC3846 digested with SphI/KpnI using NEBuilder HiFi Assembly (New England BioLabs). The *RRP40* (pAC2976) *2µ URA3* plasmid was constructed by PCR amplification of the *RRP40* gene with oligonucleotides AC4281 and AC4282 (Integrated DNA Technologies) using *S. cerevisiae* genomic DNA and cloning of BamHI/SacI-digested *RRP40* PCR product into pRS426 (92) digested with BamHI/SacI. The *MPP6* (pAC3427) *CEN6 HIS3* plasmid was constructed by PCR amplification of the *MPP6* gene with oligonucleotides AC6708 and AC7972 (Integrated DNA Technologies) using *S. cerevisiae* genomic DNA and cloning of BamHI/SacI-digested *MPP6* PCR product into pRS313 (67) digested with BamHI/SacI. The *mpp6-R112A-F115A* (pAC3433), *mpp6-R112A* (pAC3434), and *mpp6-F115A* (pAC3435) *CEN6 HIS3* plasmids were generated by site-directed mutagenesis of the *MPP6* (pAC3427) *CEN6 LEU2* plasmid using oligonucleotides encoding each amino acid change (AC7976-7981; Integrated DNA Technologies; **Table S3**) and QuickChange II SDM Kit (Agilent). The *MPP6* (pAC4561), *mpp6-R112A-F115A* (pAC4562), *mpp6-R112A* (pAC4563), and *mpp6-F115A* (pAC4564) *2µ URA3* plasmids were constructed by subcloning the *MPP6/mpp6* genes from pAC3427/3433-3435 plasmids into pRS426 (67) with BamHI/SacI digestion. The pcDNA3-2xMyc-*Exosc3* (pAC3402) plasmid was generated by PCR amplification of the mouse *Exosc3* coding sequence from Neuro-2a cDNA using oligonucleotides AC6567 and AC6568 (Integrated DNA technologies) and cloning of BamHI/XbaI-digested *Exosc3* PCR product into pcDNA3 (Invitrogen) plasmid containing pCMV promoter and N-terminal 2xMyc tag digested with BamHI/XbaI. The pcDNA3-*2xMyc-Exosc3-V27G* (pAC4526), -*2xMyc-Exosc3-G31A* (pAC3403), -*2xMyc-Exosc3-V80F* (pAC4527), -*2xMyc-Exosc3-Y108N* (pAC4528), -*2xMyc-Exosc3-D131A* (pAC3404), -*2xMyc-Exosc3-G134E* (pAC4529), -*2xMyc-Exosc3-G190C*, and -*2xMyc-Exosc3-W237R* (pAC3405) plasmids were generated by site-directed mutagenesis of pcDNA3-Myc-*Exosc3* (pAC3402) plasmid using oligonucleotides encoding each amino acid change (AC6594-6595; AC7017-7020; AC10180-10189 Integrated DNA Technologies; **Table S3**) and QuickChange II SDM Kit (Agilent). All plasmids were sequenced to ensure the presence of desired mutations and absence of any other mutations.

### Generation of *integrated* rrp40-Y64N mutant strains using CRISPR-Cas9 genome editing

Three *rrp40-Y64N* (ACY3290-3292) mutant strains were generated using CRISPR/Cas9 editing with a single pCas9-gRNA expression plasmid and double-stranded homology-directed repair (HDR) oligonucleotides in a wildtype BY4741 strain essentially as described before (91). The single pCas9-gRNA construct on a pRS316 (67) *CEN6 URA3* plasmid is derived from p414-*TEF1p-Cas9-CYC1t* plasmid (Addgene #43802) and p426-*SNR52p-gRNA.CAN1.Y-SUP4t* plasmid (Addgene #43803) (91). Constitutive expression of Cas9 is driven by the *TEF1* promoter and constitutive expression of the gRNA is driven by the *SNR52* promoter. Specifically, 500 ng of pAC3846 (pCas9 without gRNA), pAC4560 (pCas9 + *RRP40* gRNA) +/-1 nmol of double-stranded *rrp40-Y64N* HDR oligonucleotide (AC10661/10662; **Table S3**) and 50 µg salmon sperm DNA was transformed into wildtype BY4741 cells by standard Lithium Acetate transformation protocol (85). Cells were plated on Ura^-^ media plates and incubated at 30°C for 2 days. Large colonies on plates with cells transformed pCas9-gRNA and HDR oligonucleotides were restreaked to new Ura^-^ media plates and screened for the presence of *rrp40-Y64N* mutations via Sanger sequencing of genomic *RRP40* PCR products.

### *S. cerevisiae* cell growth assays

The *rrp40*Δ (yAV1107), *rrp40*Δ *mpp6*Δ (ACY2638), *rrp40*Δ *rrp47*Δ (ACY2462), and *rrp40*Δ *rrp6*Δ (ACY2466) yeast strains were transformed with wildtype *RRP40* (pAC3652) or *rrp40* mutant (*F4G/ G8A/I34F/Y64N/S87A/N90E/G148C*/*W195R*; pAC3653-3655; pAC4515-4519) *CEN6 LEU2* plasmid and selected on Leu- plates. The Leu+ transformants were streaked and grown on 5-fluoroorotic acid (5-FOA) Leu- plates to select for cells that had lost the *RRP40 CEN6 URA3* maintenance plasmid (93), and thus only contained the low copy, wildtype *RRP40* or *rrp40* mutant *CEN6 LEU2* plasmid. Cells were also transformed with empty vector (pRS315) as a control. The *rrp40*Δ *mtr4*Δ (ACY2540) yeast strain was co-transformed with *MTR4* (pAC4096), *mtr4-F7A-F10A* (pAC4099), *mtr4-R349E-N352E* (pAC4100), or *mtr4-R1030A* (pAC4104) *CEN6 HIS3* plasmid and wildtype *RRP40* (pAC3652) or *rrp40* mutant (pAC3653-3655; pAC4515-4519) *CEN6 LEU2* plasmid and selected on His- Leu- plates. The His+ Leu+ transformants were streaked and grown on 5-FOA His- Leu- plates to select for cells that had lost the wildtype *MTR4 RRP40 CEN6 URA3* maintenance plasmid (93), and thus only contained the wildtype *MTR4* or *mtr4* mutant *CEN6 HIS3* plasmid and wildtype *RRP40* or *rrp40* mutant *CEN6 LEU2* plasmid. The growth of the *rrp40*Δ, *rrp40*Δ *mpp6*Δ, *rrp40*Δ *rrp47*Δ, *rrp40*Δ *rrp6*Δ, *rrp40*Δ *MTR4*, *rrp40*Δ *mtr4-F7A-F10A*, *rrp40*Δ *mtr4-R349E-N352E*, and *rrp40*Δ *mtr4-R1030A* cells containing *RRP40* or *rrp40 CEN6 LEU2* plasmid was assayed in solid and liquid media growth assays. For growth on solid media, cells were grown to saturation in 2 ml Leu- media overnight at 30°C, serial diluted (in 10-fold dilutions), spotted on Leu- media plates, and incubated on plates at 25°C, 30°C and 37°C for 2-3 days.

The solid media growth assays presented are representative of triplicate experiments. For growth in liquid culture, saturated overnight cultures grown in Leu- media at 30°C were diluted to an A_600_ ∼ 0.01 in Leu- media in a 24-well plate, and growth at 37°C was monitored and recorded at A_600_ in a BioTek Synergy MX microplate reader with Gen5 v2.04 software over 24 hr. Technical triplicates for each biological sample were grown.

### rrp40-Y64N and rrp40-W195R CRISPR mutant cell growth assays

Wildtype (BY4741) cells, *rrp40-W195R* (ACY3117) CRISPR mutant strain, and three *rrp40-Y64N* (ACY3290-3292) CRISPR mutant strains (ACY3290-3292) were grown to saturation in 2 ml YEPD (yeast extract, peptone, dextrose) media overnight at 30°C, serial diluted (in 10-fold dilutions), spotted on YEPD media plates, and incubated on plates at 25°C, 30°C and 37°C for 2-3 days. The *rrp40-Y64N* (ACY3290-3292) CRISPR mutant strains transformed with empty vector (pRS426) or *RRP40* (pAC2976) *2µ URA3* plasmid and wildtype (BY4741) cells transformed with empty vector (pRS426) were grown in 2 ml Ura- media overnight at 30°C, serially diluted, spotted onto Ura- plates, and incubated at 25°C, 30°C and 37°C for 2-3 days as indicated. The *rrp40-W195R* (ACY3117) CRISPR mutant strain transformed with empty vector (pRS426), *RRP40* (pAC2976), or *MPP6* (pAC4561) *2µ URA3* plasmid, *rrp40-Y64N* (ACY3290) CRISPR mutant strain transformed with empty vector (pRS426), *RRP40* (pAC2976), *MPP6* (pAC4561), *mpp6-R112A-F115A* (pAC4562), *mpp6-R112A* (pAC4563), or *mpp6-F115A* (pAC4564) *2µ URA3* plasmid, and wildtype (BY4741) cells transformed with empty vector (pRS426) were grown in 2 ml Ura- media overnight at 30°C, serially diluted, spotted onto Ura- plates, and incubated at 25°C, 30°C and 37°C for 2-3 days as indicated. The *rrp40-W195R mpp6*Δ (ACY3321) strain and *rrp40-Y64N mpp6*Δ (ACY3327) strain transformed with empty vector (pRS313), *MPP6* (pAC3427), *mpp6-R112A-F115A* (pAC3433), *mpp6-R112A* (pAC3434), or *mpp6- F115A* (pAC3435) *CEN6 HIS3* plasmid, and wildtype (BY4741), *rrp40-W195R* (ACY3117) CRISPR mutant, *rrp40-Y64N* (ACY3290) CRISPR mutant, and *mpp6*Δ (ACY3315) cells transformed with empty vector (pRS313) were grown in 2 ml His- media overnight at 30°C, serially diluted, spotted onto His- plates, and incubated at 25°C, 30°C and 37°C for 2-4 days as indicated.

### Model building

To construct a model of the budding yeast RNA exosome complex, we used the structures of the yeast RNA exosome core (PDB ID: 6FSZ) (78) and the Mpp6–nuclear RNA exosome complex bound to RNA (PDB ID: 5VZJ) (76) as templates. We first replaced the active Dis3 subunit in 6FSZ with the corresponding subunit from 5VZJ. The unresolved regions of Rrp6 (residues 106-149, 400-538, and 557-565), together with missing regions in other subunits, were modeled using AlphaFold2 predictions. The full RNA strand was generated by joining the RNA fragments from 5VZJ and 6FSZ.

To construct a model of the human RNA exosome complex, we began with the Cryo-EM structure of the human nuclear exosome–MTR4–RNA complex (PDB ID: 6D6R) (83) and its corresponding density map (EMDB accession code: EMD-22587). The RNA exosome cofactor MTR4 and the RNA exosome component EXOSC10 were positioned within the RNA exosome based on structural predictions from AlphaFold2 (79,94) and AlphaFold3-multimer (95). First, we generated MTR4/EXOSC10 complexes together with EXOSC2/EXOSC3/EXOSC4/EXOSC7/EXOSC9/MPP6/RNA using AlphaFold3-multimer. The MTR4 (residues 596-843)/EXOSC10 (residues 21-591) module was placed by aligning it to the 6D6R structure. The remaining region of EXOSC10 (residues 592–804) was positioned according to the AlphaFold2 prediction. To assemble the complete human RNA exosome complex, we additionally modeled unresolved loop regions of the core subunits (EXOSC1–10, MPP6, MTR4, and DIS3) using AlphaFold2 predictions.

Following construction of both the yeast and human RNA exosome complex models, each structure was subjected to energy minimization in Phenix (96) to relieve local steric clashes and improve geometric consistency introduced during model assembly. The resulting minimized models were evaluated for stereochemical quality and overall structural validity using MolProbity (97), with summary validation statistics, including Ramachandran outliers, rotamer outliers, and clashscore, provided in **Table S4**. Both the budding yeast and human RNA exosome complex models have been deposited in ModelArchive.

### Rosetta Protein Stability Analysis

To assess the impact of disease-associated amino acid substitutions in EXOSC3, protein stability was predicted using the Rosetta Cartesian ΔΔG protocol (98). The wildtype structure of the RNA exosome complex was first optimized in Cartesian space using the Rosetta Fast Relax protocol. Amino acid substitutions were then introduced, followed by repacking of side chains within 6Å of the amino acid substitution using the FastRelax protocol. Additionally, the protein backbone was allowed to readjust within three residues of the mutation site. The ΔΔG values were calculated as the difference in Rosetta scores between the relaxed mutant and relaxed wildtype proteins for each analyzed amino acid substitution.

### Immunoblotting

For analysis of C-terminally Myc-tagged wildtype Rrp40 and rrp40 variant protein expression levels, *rrp40*Δ cells (yAV1107) were transformed with wildtype *RRP40-2xMyc* (pAC3666) or *rrp40-2xMyc* mutant (*F4G/G8A/I34F/Y64N/S87A/N90E/G148C/W195R*; pAC3667-3668; pAC4520-4525) *CEN6 LEU2* plasmid and selected on Leu- plates. The Leu+ transformants were streaked and grown on 5-FOA

Leu- plates to select for cells that had lost the *RRP40 CEN6 URA3* maintenance plasmid (93), and thus only contained the low copy, wildtype *RRP40-2xMyc* or *rrp40-2xMyc* mutant *CEN6 LEU2* plasmid. Cells were also transformed with empty vector (pRS315) as a control. The *rrp40*Δ cells only expressing wildtype Rrp40-2xMyc or rrp40-2xMyc variants were grown in 2 ml Leu- media overnight at 30°C to saturation and 10 ml cultures with an A_600_ = 0.3 were prepared and incubated at 30 or 37°C for 5 hr. Cell pellets were collected by centrifugation, transferred to 2 ml screw-cap tubes and stored at −80°C. For analysis of N-terminally Myc-tagged EXOSC3 expression levels, mouse Neuro-2a (N2a) cells (99) were transiently transfected with pcDNA3 vector (Invitrogen) containing mouse, wildtype 2x*Myc-Exosc3* (pAC3402) or 2x*Myc-Exosc3* variant (*V27G/G31A/V80F/Y108N/D131A/G134E/G190C/W237R*; pAC3403-3405; pAC4526-4530), or empty vector (pcDNA3; Invitrogen) using Lipofectamine 2000 (Invitrogen) and cells were collected 24 hr after transfection.

Budding yeast cell lysates were prepared by resuspension of cells in 0.5 ml RIPA-2 Buffer [50 mM Tris-HCl, pH 8; 150 mM NaCl; 0.5% sodium deoxycholate; 1% NP40; 0.1% SDS] supplemented with protease inhibitors [1 mM PMSF; Pierce™ Protease Inhibitors (Thermo Fisher Scientific)], addition of 300 µl glass beads, disruption in a Mini Bead Beater 16 Cell Disrupter (Biospec) for 4 x 1 min at 25°C, and centrifugation at 16,000*g* for 20 min at 4°C. Mouse N2a cell lysates were prepared by lysis in RIPA-2 Buffer and centrifugation at 16,000*g* for 10 min at 4°C. Protein lysate concentration was determined by Pierce BCA Protein Assay Kit (Life Technologies). Whole cell lysate protein samples (20-50 µg) were resolved on Criterion 4-20% gradient denaturing gels (Bio-Rad), transferred to nitrocellulose membranes (Bio-Rad) and Myc-tagged Rrp40, and EXOSC3 proteins were detected with anti-Myc monoclonal antibody 9B11 (1:2000; Cell Signaling; Cat. 2276S). As loading controls, 3- Phosphoglycerate kinase (Pgk1) protein was detected with anti-Pgk1 monoclonal antibody (1:30,000; Invitrogen; Cat. 459250) and Stain-Free signal on the immunoblot was included. For transfection control, Neomycin phosphotransferase II (NPTII) expressed from *NeoR* cassette on pcDNA3*-Exosc3* plasmids was detected with anti-NPTII monoclonal antibody (1:1000; Cell Applications, Inc.; Cat. CP10330). As a loading control, Beta-actin (ACTB) was detected with anti-ACTB monoclonal antibody (1:2000; Proteintech; Cat. 66009-1-Ig). Primary antibodies were detected using goat anti-mouse secondary antibodies coupled to horseradish peroxidase (1:3000; Jackson ImmunoResearch Inc; Cat. 115-035-003) and enhanced chemiluminescence signals were captured on a ChemiDoc Imaging System (Bio-Rad).

### Quantitation of immunoblotting

The protein band intensities/areas from immunoblots were quantitated using ImageJ v1.4 software (National Institute of Health, MD; http://rsb.info.nih.gov/ij/) or ImageLab software (Bio-Rad) and mean fold changes in protein were calculated in Microsoft Excel for Mac Version 16.110.3 (Office 365, Microsoft Corporation). To quantitate the levels of rrp40-2xMyc variants relative to wildtype Rrp40- 2xMyc in *rrp40*Δ cells incubated at 30°C or 37°C, wildtype Rrp40-2xMyc and rrp40-2xMyc variant intensity was first normalized to loading control Pgk1 intensity and then normalized to wildtype Rrp40- 2xMyc intensity at 30°C or 37°C. To calculate the levels of 2xMyc-EXOSC3 variants relative to wildtype 2xMyc-EXOSC3 in N2a cells, wildtype 2xMyc-EXOSC3 and 2xMyc-EXOSC3 variant intensity were first normalized to transfection control NPTII intensity, then normalized to loading control ACTB intensity, and finally normalized to wildtype Myc-EXOSC3 intensity. The level of rrp40- 2xMyc variants relative to wildtype Rrp40-2xMyc and 2xMyc-EXOSC3 variants relative to wildtype 2xMyc-EXOSC3 were calculated and values are indicated below each blot. Results shown are representative of independent experiments.

### Total RNA isolation from S. cerevisiae

To prepare *S. cerevisiae* total RNA, cells were grown in 10 ml cultures to A_600_ = 0.5-0.8 at 30°C. Cell pellets in 2 ml screw-cap tubes were resuspended in 1 ml TRIzol (Invitrogen), 300 µl glass beads were added, and samples were disrupted in a Mini Bead Beater 16 Cell Disrupter (Biospec) for 2 min at 25°C. For each sample, 100 µl of 1-bromo-3-chloropropane (BCP) was added, the sample was vortexed for 15 sec, and incubated at 25°C for 2 min. Each sample was centrifuged at 16,300*g* for 8 min at 4°C and upper layer was transferred to a fresh microfuge tube. RNA was precipitated with 500 µl isopropanol and sample was vortexed for 10 sec to mix. Total RNA was pelleted by centrifugation at 16,300*g* for 8 min at 4°C. RNA pellet was washed with 1 ml of 75% ethanol, centrifuged at 16,300*g* for 5 min at 4°C, and air dried for 15 min. Total RNA was resuspended in 50 µl diethylpyrocarbonate (DEPC (Sigma))- treated water and stored at −80°C.

### Quantitative RT-PCR

For analysis of *U4* pre-snRNA, *TLC1* pre-RNA, and *U14* snoRNA levels in wildtype *RRP40* and *rrp40* mutant cells, *rrp40*Δ (yAV1107) cells solely containing *RRP40* (pAC3652), *rrp40-W195R* (pAC3655), or *rrp40-Y64N* (pAC4517) *CEN6 LEU2* plasmid were grown in biological triplicate in 2 ml Leu- media overnight at 30°C, 10 ml cultures with an A_600_ = 0.4 were prepared and grown at 37°C for 5 hr. Cells were collected by centrifugation (2,163*g*; 16,000*g*), transferred to 2 ml screw cap tubes and stored at −80°C. Following total RNA isolation from each cell pellet, 1 µg RNA was reverse transcribed to 1^st^ strand cDNA using the M-MLV Reverse Transcriptase (Invitrogen) and 0.3 µg random hexamers according to manufacturer’s protocol. Quantitative PCR was performed on technical triplicates of cDNA (10 ng) from independent biological triplicates using gene specific primers to detect *U4* pre-snRNA, *TLC1* pre-RNA, and *U14* snoRNA (0.5 µM; AC5722-5723; AC7593-7594; AC5397-5398; **Table S3**) and QuantiTect SYBR Green PCR master mix (Qiagen) on a StepOnePlus Real-Time PCR machine (Applied Biosystems; T_anneal_ = 55°C; 44 cycles). Primers to detect *ALG9* mRNA were used as a normalization control (AC5067-5068; **Table S3**). The mean RNA levels were calculated by the ΔΔCt method (100), normalized to mean RNA levels in *RRP40* cells, and converted and graphed as RNA fold change relative to *RRP40* with error bars that represent the standard error of the mean.

### Northern blot analysis of rRNAs

For the analysis of ribosomal RNAs, triplicate samples of wildtype (BY4741) cells, three *rrp40-W195R* (ACY3117-3119) CRISPR mutant strains, and three *rrp40-Y64N* (ACY3290-3292) CRISPR mutant strains were inoculated into 2 ml yeast extract, peptone, dextrose (YEPD) media and grown at 30°C overnight to saturation. The overnight cultures were rediluted in 10 ml YEPD media to an A_600_ = 0.1 for day cultures and grown to log phase at 30°C to an A_600_ = 0.4 to 0.6. The log phase, day cultures were rediluted in 10 ml YEPD media for final cultures to an A_600_ = 0.00025 to 0.004 and grown overnight to log phase at 30°C or 37°C to an A_600_ = 0.5-0.6. The final cultures (10 ml) were collected by centrifugation at 1962*g* for 3 min, transferred in 1 ml water to 2 ml screw-cap tubes, centrifuged at 16,000*g* for 1 min, aspirated, and cell pellets were frozen in liquid nitrogen, and stored at −80°C. Total RNA was extracted using the hot phenol method (101). Northern blotting was carried out essentially as previously described (102) using oligonucleotide probes for 7S (ITS2) rRNA and loading control 5S rRNA (**Table S3**).

### Cell Culture

Neuro-2a (N2a) cells are a mouse neuroblastoma cell line (103). Cells were maintained in Dulbecco’s modified Eagle’s medium (DMEM) supplemented with 10% FBS and antibiotics and grown under standard environmental conditions. The authenticity of the cell line was validated using STR profiling. In addition, the cell line was tested for mycoplasma, and cells were free from mycoplasma contamination for all experiments.

### Immunoprecipitation of mammalian RNA exosome subunits

To immunoprecipitate mouse EXOSC3, murine Neuro-2a cells (N2a) were transiently transfected with 2x*Myc-Exosc3* (pAC3402) or *2xMyc-Exosc3* variants (pAC3403-3405; pAC4526-4530) using Lipofectamine 2000 (Invitrogen) and cells were collected 48 hr after transfection. Cells were lysed in HEPES binding buffer [20 mM HEPES, pH 7.2; 150 mM NaCl; 0.2% Triton X-100; Pierce™ Protease Inhibitors (Thermo Fisher Scientific)]. Cell lysates (250-300 µg) in 0.3 ml HEPES binding buffer were incubated with 20 µl Pierce™ anti-c-Myc magnetic beads (Thermo Fisher Scientific) for 3 hr at 4°C with mixing. Beads were pelleted by centrifugation, unbound supernatant was removed, and beads were washed three times with 0.3 ml binding buffer. Input (25 µg) and Bound Myc bead (1/2 total amount) samples were analyzed by SDS-PAGE and immunoblotting with mouse anti-Myc monoclonal antibody (1:2000; Cell Signaling; Cat. 2276S) to detect Myc-tagged EXOSC3, rabbit anti-EXOSC8 (1:1000; Proteintech; Cat. 11979-1-AP) to detect endogenous EXOSC8, rabbit anti-EXOSC9 (1:1000; Bethyl Laboratories, Inc.; Cat. A303-888A) to detect endogenous EXOSC9, or rabbit anti-EXOSC10 (1:1000; Bethyl Laboratories, Inc.; Cat. A303-987A) to detect endogenous EXOSC10. Primary antibodies were detected using goat anti-mouse or goat anti-rabbit secondary antibodies coupled to horseradish peroxidase (1:3000; Jackson ImmunoResearch Inc; Cat. 115-035-003 and Cat. 111-035-003) and enhanced chemiluminescence signals were captured on a ChemiDoc Imaging System (Bio-Rad).

## Data Availability Statement

Strains and plasmids are available upon request. The models are available in ModelArchive (modelarchive.org): ma-k3wui (*S. cerevisiae*, access code: uVLWa8BGte) and ma-64tn4 (human, access code: gOlx3S5nOM). The authors affirm that all data necessary for confirming the conclusions of the article are present within the article, figures, and tables.

## ACKNOWLEDGEMENTS

We thank members of the research groups for project discussions and experimental insight. This study was supported by NIH awards 2R35GM138123-06 to H.G, R21AI193385 to AHC, and R35GM139382 and R01ES032786 to I.I. M.L.I. was supported by NIH T32GM135060. An award of computer time to I.I. was provided by the INCITE program. This research also used resources of the Oak Ridge Leadership Computing Facility, which is a DOE Office of Science User Facility supported under Contract [DE-AC05-00OR22725].

