## Supplementary material for "Systematic comparison of pathogenic variants of the RNA exosome gene *EXOSC3/RRP40* in *Saccharomyces cerevisiae* reveals variant-specific functional consequences": Table S1


Figure S1. Immunoblot to detect Rrp40 and rrp40 variants including total protein loading control..S-2

Figure S2. The *rrp40-Y64N* mutant cells exhibit defects in RNA processing....................................S-3

Figure S3. Overexpression of Mpp6 rescues growth defects of *rrp40-Y64N* mutant cells.................S-4

Table S1: Disease-associated EXOSC3 variants……………………………………………………….S-5

Table S2. *Saccharomyces cerevisiae* strains and plasmids used in this study...................................S-6

Table S3. DNA oligonucleotide primers and probes used in this study............................................S-8

Table S4. MolProbity results for budding yeast and human RNA exosome models……………….S-10

Supporting References...........................................................................………………………S-11

**Figure S1**

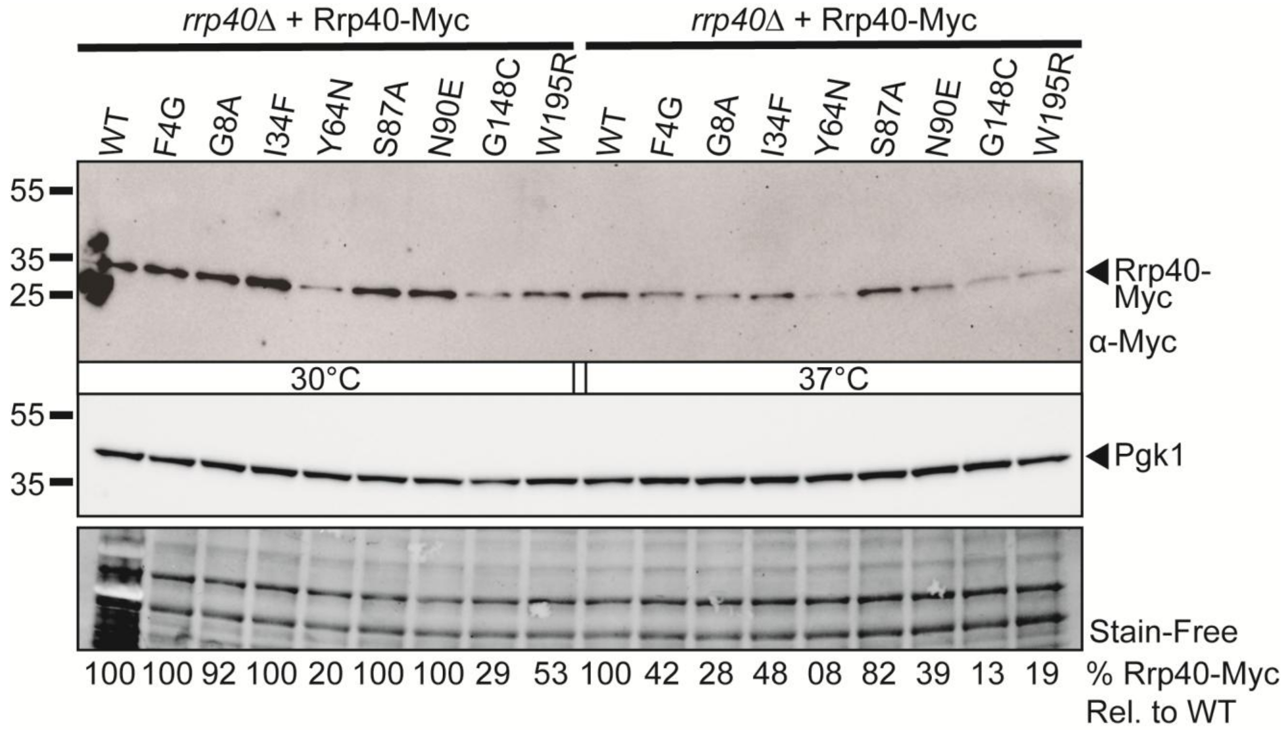


**Figure S1 Immunoblot to detect Rrp40 and rrp40 variants including total protein loading control.** The steady-state level of some Rrp40 protein variants is decreased at 30°C and 37°C. Lysates of *rrp40Δ* cells solely expressing Myc-tagged, wildtype Rrp40 (WT) or rrp40 variants grown at 30°C or 37°C were analyzed by immunoblotting with an anti-Myc antibody to detect Rrp40-Myc and an anti-Pgk1 antibody to detect 3-phosphoglycerate kinase (Pgk1) as a loading control. As an additional loading control total protein was detected with Stain-Free. Quantitation of the percentage of the rrp40 protein level relative to Rrp40 is indicated below with WT Rrp40 at each temperature set to 100%. Results are representative of two independent experiments.

**Figure S2**

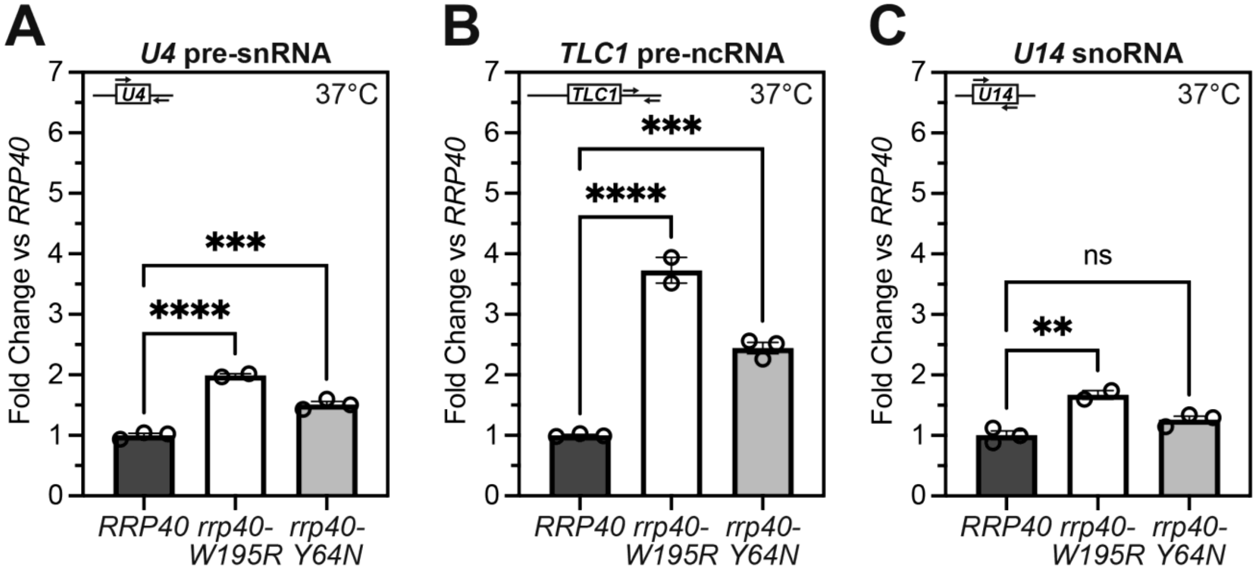


**Figure S2 The *rrp40-Y64N* mutant cells exhibit defects in RNA processing.** RT-qPCR analysis of RNA exosome target RNAs reveals RNA processing defects in *rrp40-Y64N* mutant cells. Total RNA was isolated from *rrp40*Δ cells solely expressing *RRP40*, *rrp40-W195R* or *rrp40-Y64N* cells grown at 37°C and transcript levels were measured by RT-qPCR using gene specific primers, normalized relative to *RRP40* for: *A*, *U4* pre-snRNA, **B**, *TLC1* telomerase component pre-RNA, and *C*, *U14* snoRNA relative to *RRP40* cells at 37°C. Error bars represent standard error of the mean from three biological replicates. Statistical significance of the RNA levels in *rrp41-L187P* cells relative to *RRP41* cells was calculated by t-test (***P-value ≤ 0.001; ****P-value ≤ 0.0001).

**Figure S3**

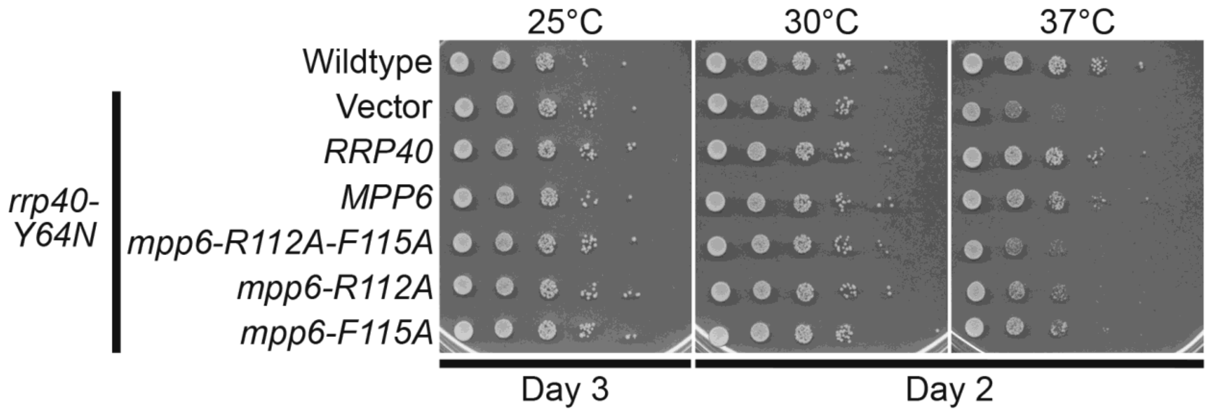


**Figure S3 Overexpression of Mpp6 rescues growth defects of *rrp40-Y64N* mutant cells.** Plasmids that overexpress *RRP40*, *MPP6*, *mpp6-R112A-F115A*, *mpp6-R112A*, or *mpp6-F115A* as well as a Vector control were transformed into *rrp40-Y64N* mutant cells and growth at the indicated temperatures was assayed by serial dilution growth assay. Wildtype cells are also included as a control. Mpp6 R112 and F115 are key residues required for functional Mpp6 interaction with Rrp40 and stimulation of RNA exosome activity (1,2). These transformed cells as well as Wildtype control cells were analyzed by serial dilution growth assay for the indicated number of Days at 25°C, 30°C or 37°C. Results shown are representative of at least three independent experiments.

**Table S1: Disease-associated EXOSC3 variants**

| RNA Exosome Subunit | Amino Acid Substitution | Genotype | Zygosity | Patient Diagnosis# | Age of Onset of Symptoms* |
|---|---|---|---|---|---|
| EXOSC3 | V27G | V27G/Null | Compound heterozygous | PCH1b | Childhood to adolescence |
| EXOSC3 | G31A | G31A | Homozygous | PCH1b | Infancy |
| EXOSC3 | V80F | Y80F/D132A | Compound heterozygous | PCH1b, CSP | Childhood |
| EXOSC3 | Y109N | Y109N/D132A | Compound heterozygous | PCH1b | Infancy (neonatal to early infancy) |
| EXOSC3 | D132A | D132A | Homozygous | PCH1b | Infancy |
| | | D132A/Null | Compound heterozygous | PCH1b | Infancy to adolescence |
| EXOSC3 | G135E | G135E | Homozygous | PCH1b | Infancy |
| EXOSC3 | G191C | G191C | Homozygous | PCH1b, CSP | Childhood to adolescence |
| EXOSC3 | W238R | W238R/G31A | Compound heterozygous | PCH1b | Infancy |

*Present at Birth, neonatal (1st month), infancy (1-12 months), toddler (12-24 months), early childhood (2-5 years), middle childhood (6-11 years), adolescents (11-12 years);
#PCH1b, Pontocerebellar Hypoplasia Type 1b; CSP, Complicated spastic paraplegia

**Table S2.** ***Saccharomyces cerevisiae*** **strains and plasmids used in this study**

| Strain/Plasmid | Description | Reference |
|---|---|---|
| BY4741 (ACY402) | *MATa*, *ura3Δ0*, *leu2Δ0*, *his3Δ1*, *met15Δ0* | |
| *rrp40Δ* (yAV1107) | *MATa*, *ura3Δ0*, *leu2Δ0*, *his3Δ1*, *RRP40::neoMX*, [*RRP40*, *URA3*] | (3) |
| *rrp40Δ mpp6Δ* (ACY2638) | *MATa*, *ura3Δ0*, *leu2Δ0*, *his3Δ1*, *RRP40::neoMX*, *MPP6::natMX4*, [*RRP40*, *URA3*] | (4) |
| *rrp40Δ rrp47Δ* (ACY2462) | *MATa*, *ura3Δ0*, *leu2Δ0*, *his3Δ1*, *RRP40::neoMX*, *LRP1::natMX4*, [*RRP40*, *URA3*] | (4) |
| *rrp40Δ rrp6Δ* (ACY2466) | *MATa*, *ura3Δ0*, *leu2Δ0*, *his3Δ1*, *RRP40::neoMX*, *RRP6::natMX4*, [*RRP40*, *URA3*] | (4) |
| *rrp40Δ mtr4Δ* (ACY2540) | *MATalpha, ura3-1, leu2-3,112, his3-11,15, trp1-1, RRP40::neoMX, MTR4::NatMX4,* [*RRP40*, *MTR4*, *URA3*] | (4) |
| *rrp40-W195R* (1) (ACY3117) | *MATa*, *ura3Δ0*, *leu2Δ0*, *his3Δ1*, *met15Δ0*, *rrp40-W195R* | (5) |
| *rrp40-W195R* (2) (ACY3118) | *MATa*, *ura3Δ0*, *leu2Δ0*, *his3Δ1*, *met15Δ0*, *rrp40-W195R* | This Study |
| *rrp40-W195R* (3) (ACY3119) | *MATa*, *ura3Δ0*, *leu2Δ0*, *his3Δ1*, *met15Δ0*, *rrp40-W195R* | This Study |
| *rrp40-Y64N* (1) (ACY3290) | *MATa*, *ura3Δ0*, *leu2Δ0*, *his3Δ1*, *met15Δ0*, *rrp40-Y64N* | This Study |
| *rrp40-Y64N* (2) (ACY3291) | *MATa*, *ura3Δ0*, *leu2Δ0*, *his3Δ1*, *met15Δ0*, *rrp40-Y64N* | This Study |
| *rrp40-Y64N* (3) (ACY3292) | *MATa*, *ura3Δ0*, *leu2Δ0*, *his3Δ1*, *met15Δ0*, *rrp40-Y64N* | This Study |
| *mpp6Δ* (ACY3315) | *MATa*, *ura3Δ0*, *leu2Δ0*, *his3Δ1*, *met15Δ*, *MPP6::natMX4* | This Study |
| *rrp40-W195R mpp6Δ* (ACY3321) | *MATa*, *ura3Δ0*, *leu2Δ0*, *his3Δ1*, *met15Δ0*, *rrp40-W195R, MPP6::natMX4* | This Study |
| *rrp40-Y64N mpp6Δ* (ACY3327) | *MATa*, *ura3Δ0*, *leu2Δ0*, *his3Δ1*, *met15Δ0*, *rrp40-Y64N, MPP6::natMX4* | This Study |
| pRS313 (pAC1) | *CEN6*, *HIS3*, $amp^R$ | (6) |
| pRS315 (pAC3) | *CEN6*, *LEU2*, $amp^R$ | (6) |
| pRS426 (pAC8) | *2µ, URA3,* $amp^R$ | (7) |
| pAC3652 | *RRP40* in pRS315, *CEN6*, *LEU2*, $amp^R$ | (4) |
| pAC3653 | *rrp40-G8A* in pRS315, *CEN6*, *LEU2*, $amp^R$ | This Study |
| pAC3654 | *rrp40-S87A* in pRS315, *CEN6*, *LEU2*, $amp^R$ | (5) |
| pAC3655 | *rrp40-W195R* in pRS315, *CEN6*, *LEU2*, $amp^R$ | (4) |
| pAC4515 | *rrp40-F4G* in pRS315, *CEN6*, *LEU2*, $amp^R$ | This Study |
| pAC4516 | *rrp40-I34F* in pRS315, *CEN6*, *LEU2*, $amp^R$ | This Study |
| pAC4517 | *rrp40-Y64N* in pRS315, *CEN6*, *LEU2*, $amp^R$ | This Study |
| pAC4518 | *rrp40-N90E* in pRS315, *CEN6*, *LEU2*, $amp^R$ | This Study |
| pAC4519 | *rrp40-G148C* in pRS315, *CEN6*, *LEU2*, $amp^R$ | This Study |
| pAC3666 | *RRP40-2xMyc* in pRS315, *CEN6*, *LEU2*, $amp^R$ | This Study |

| | | |
|---|---|---|
| pAC4521 | *rrp40-F4G-2xMyc* in pRS315, *CEN6*, *LEU2*, *amp*$^R$ | This Study |
| pAC4520 | *rrp40-G8A-2xMyc* in pRS315, *CEN6*, *LEU2*, *amp*$^R$ | This Study |
| pAC4522 | *rrp40-I34F-2xMyc* in pRS315, *CEN6*, *LEU2*, *amp*$^R$ | This Study |
| pAC4523 | *rrp40-Y64N-2xMyc* in pRS315, *CEN6*, *LEU2*, *amp*$^R$ | This Study |
| pAC3667 | *rrp40-S87A-2xMyc* in pRS315, *CEN6*, *LEU2*, *amp*$^R$ | This Study |
| pAC4524 | *rrp40-N90E-2xMyc* in pRS315, *CEN6*, *LEU2*, *amp*$^R$ | This Study |
| pAC4525 | *rrp40-G148C-2xMyc* in pRS315, *CEN6*, *LEU2*, *amp*$^R$ | This Study |
| pAC3668 | *rrp40-W195R-2xMyc* in pRS315, *CEN6*, *LEU2*, *amp*$^R$ | This Study |
| pAC4096 | *MTR4* in pRS313, *CEN6*, *HIS3*, *amp*$^R$ | (4) |
| pAC4099 | *mtr4-F7A-F10A* in pRS313, *CEN6*, *HIS3*, *amp*$^R$ | (4) |
| pAC4100 | *mtr4-R349E-N352E* in pRS313, *CEN6*, *HIS3*, *amp*$^R$ | (8) |
| pAC4104 | *mtr4-R1030A* in pRS313, *CEN6*, *HIS3*, *amp*$^R$ | (8) |
| pAC3846 | *TEF1p-Cas9-CYC1t-SNR52p* in pRS316, *CEN6*, *URA3*, *amp*$^R$ | (5) |
| pAC4560 | *TEF1p-Cas9-CYC1t-SNR52p-RRP40_164.gRNA-SUP4t* in pRS316, *CEN6*, *URA3*, *amp*$^R$ | This Study |
| p4339 (pAC1992) | *TA::natMX4* switcher cassette in pCRII-TOPO*, amp*$^R$ | (9) |
| pAC2976 | *RRP40* in pRS426, *2μ*, *URA3*, *amp*$^R$ | This Study |
| pAC3427 | *MPP6* in pRS313, *CEN6*, *HIS3*, *amp*$^R$ | This Study |
| pAC3433 | *mpp6-R112A-F115A* in pRS313, *CEN6*, *HIS3*, *amp*$^R$ | This Study |
| pAC3434 | *mpp6-R112A* in pRS313, *CEN6*, *HIS3*, *amp*$^R$ | This Study |
| pAC3435 | *mpp6-F115A* in pRS313, *CEN6*, *HIS3*, *amp*$^R$ | This Study |
| pAC4561 | *MPP6* in pRS426, *2μ*, *URA3*, *amp*$^R$ | This Study |
| pAC4562 | *mpp6-R112A-F115A* in pRS426, *2μ*, *URA3*, *amp*$^R$ | This Study |
| pAC4563 | *mpp6-R112A* in pRS426, *2μ*, *URA3*, *amp*$^R$ | This Study |
| pAC4564 | *mpp6-F115A* in pRS426, *2μ*, *URA3*, *amp*$^R$ | This Study |
| pcDNA3 (pAC2319) | *pCMV, Neo*$^R$*, amp*$^R$ | Invitrogen |
| pAC3402 | *2xMyc-Exosc3* in pcDNA3, *Neo*$^R$, amp$^R$ | This Study |
| pAC4526 | *2xMyc-Exosc3-V27G* in pcDNA3, *Neo*$^R$, *amp*$^R$ | This Study |
| pAC3403 | *2xMyc-Exosc3-G31A* in pcDNA3, *Neo*$^R$, *amp*$^R$ | This Study |
| pAC4527 | *2xMyc-Exosc3-V80F* in pcDNA3, *Neo*$^R$, *amp*$^R$ | This Study |
| pAC4528 | *2xMyc-Exosc3-Y108N* in pcDNA3, *Neo*$^R$, *amp*$^R$ | This Study |
| pAC3404 | *2xMyc-Exosc3-D131A* in pcDNA3, *Neo*$^R$, *amp*$^R$ | This Study |
| pAC4529 | *2xMyc-Exosc3-G134E* in pcDNA3, *Neo*$^R$, *amp*$^R$ | This Study |
| pAC4530 | *2xMyc-Exosc3-G190C* in pcDNA3, *Neo*$^R$, *amp*$^R$ | This Study |
| pAC3405 | *2xMyc-Exosc3-W237R* in pcDNA3, *Neo*$^R$, *amp*$^R$ | This Study |

**Table S3. DNA oligonucleotide primers and probes used in this study**

| Description | Sequence (5'-3') | Name |
|---|---|---|
| *rrp40-F4G* Fwd | GCATACAAGATGTCTACG**GGC**ATATTCCCTGGTGATAGC | AC10170 |
| *rrp40-F4G* Rev | GCTATCACCAGGGAATAT**GCC**CGTAGACATCTTGTATGC | AC10171 |
| *rrp40-G8A* Fwd | ATGTCTACGTTCATATTCCCT**GCT**GATAGCTTTCCTGTAG | AC5187 |
| *rrp40-G8A* Rev | CTACAGGAAAGCTATC**AGC**AGGGAATATGAACGTAGACAT | AC5188 |
| *rrp40-I34F* Fwd | CCCCAATACTCAAGAA**TTT**CGACCTGTTAATACAGGTG | AC10172 |
| *rrp40-I34F* Rev | CACCTGTATTAACAGGTCG**AAA**TTCTTGAGTATTGGGG | AC10173 |
| *rrp40-Y64N* Fwd | CTATTCTAGTAAGAGA**AAC**ATTCCATCTGTAAACG | AC10174 |
| *rrp40-Y64N* Rev | CGTTTACAGATGGAAT**GTT**TCTCTTACTAGAATAG | AC10175 |
| *rrp40-S87A* Fwd | CAGATAGCTATAAGGTT**GCG**TTGCAAAATTTCTCCTCC | AC5984 |
| *rrp40-S87A* Rev | GGAGGAGAAATTTTGCAA**CGC**AACCTTATAGCTATCTG | AC5985 |
| *rrp40-N90E* Fwd | GCTATAAGGTTTCGTTGCAA**GAA**TTCTCCTCCAGTGTTTCAC | AC10176 |
| *rrp40-N90E* Rev | GTGAAACACTGGAGGAGAA**TTC**TTGCAACGAAACCTTATAGC | AC10177 |
| *rrp40-G148C* Fwd | GGACGCGATGCTGGTTTC**TGT**ATATTGGAAGATGGTATGATC | AC10178 |
| *rrp40-G148C* Rev | GATCATACCATCTTCCAATAT**ACA**GAAACCAGCATCGCGTCC | AC10179 |
| *rrp40-W195R* Fwd | GGTCTCAATGGGAAGATC**CGG**GTTAAGTGCGAGG | AC5982 |
| *rrp40-W195R* Rev | CCTCGCACTTAAC**CCG**GATCTTCCCATTGAGACC | AC5983 |
| *Exosc3* CDS Fwd | ATATGGATCCATGGCTGAAGTACTGTCCGCG | AC6567 |
| *Exosc3* CDS Rev | ATATTCTAGATCAACTCTCTGCCAGTCTGGC | AC6568 |
| *Exosc3-V27G* Fwd | GCACAAGGTGCTGAATCAG**GGG**GTTCTCCCCGGGGAGGAGC | AC10180 |
| *Exosc3-V27G* Rev | GCTCCTCCCCGGGGAGAAC**CCC**CTGATTCAGCACCTTGTGC | AC10181 |
| *Exosc3-G31A* Fwd | GTGGTTCTCCCC**GCG**GAGGAGCTGGTGCTG | AC7019 |
| *Exosc3-G31A* Rev | CAGCACCAGCTCCTC**CGC**GGGGAGAACCAC | AC7020 |
| *Exosc3-V80F* Fwd | GCTGCGGGGACCGTCTGCTG**TTC**ACCAAGTGTGGCCGCCTGC | AC10182 |
| *Exosc3-V80F* Rev | GCAGGCGGCCACACTTGGT**GAA**CAGCAGACGGTCCCCGCAGC | AC10183 |
| *Exosc3-Y108N* Fwd | GGACTCGCAGCAGAAGCGG**AAT**GTACCTGTGAAAGGGGACC | AC10184 |
| *Exosc3-Y108N* Rev | GGTCCCCTTTCACAGGTAC**ATT**CCGCTTCTGCTGCGAGTCC | AC10185 |
| *Exosc3-D131A* Fwd | GAGATATATTCAAAGTT**GCT**GTTGGAGGGAGTGAGCC | AC7017 |
| *Exosc3-D131A* Rev | GGCTCACTCCCTCCAAC**AGC**AACTTTGAATATATCTC | AC7018 |
| *Exosc3-G134E* Fwd | GATATATTCAAAGTTGATGTTGGA**GAG**AGTGAGCCAGCGTCT TTGTC | AC10186 |
| *Exosc3-G134E* Rev | GACAAAGACGCTGGCTCACT**CTC**TCCAACATCAACTTTGAAT ATATC | AC10187 |
| *Exosc3-G190C* Fwd | GGCCGCGCCAATGGGATG**TGT**GTGATTGGGCAGGATGGCC | AC10188 |
| *Exosc3-G190C* Rev | GGCCATCCTGCCCAATCAC**ACA**CATCCCATTGGCGCGGCC | AC10189 |
| *Exosc3-W237R* Fwd | GTTTGGAATGAATGGAAGAATA**AGG**GTCAAAGCTAAGACCAT TCAG | AC6594 |
| *Exosc3-W237R* Rev | CTGAATGGTCTTAGCTTTGAC**CCT**TATTCTTCCATTCATTCCAA AC | AC6595 |
| *RRP40* gene Fwd | CGGGGATCCTTGATCTGGTTTGCCTAATCC | AC4281 |
| *RRP40* gene Rev | CCGGAGCTCCGGTCATGATATTGTTGACTTTG | AC4282 |
| *MPP6* gene Fwd | ATATGGATCCCGAATCTACACGAGAGCAACAAACGATACC | AC6708 |
| *MPP6* gene Rev | ATATGAGCTCACTTACATCAAACAACTTTGTTTGGCAACC | AC7972 |

| | | |
|---|---|---|
| *mpp6-R112A-F115A* Fwd | CCTGAGGGTGTGATAAGTGGG**GCA**AAAACC**GCT**GGCGATAATTCTGATGATAGTGG | AC7976 |
| *mpp6-R112A-F115A* Rev | CCACTATCATCAGAATTATCGCC**AGC**GGTTTT**TGC**CCCACTTATCACACCCTCAGG | AC7977 |
| *mpp6-R112A* Fwd | CCTGAGGGTGTGATAAGTGGG**GCA**AAAACCTTTGGCGATAATTCTGATGATAGTGG | AC7978 |
| *mpp6-R112A* Rev | CCACTATCATCAGAATTATCGCCAAAGGTTTT**TGC**CCCACTTATCACACCCTCAGG | AC7979 |
| *mpp6-F115A* Fwd | CCTGAGGGTGTGATAAGTGGGAGAAAAACC**GCT**GGCGATAATTCTGATGATAGTGG | AC7980 |
| *mpp6-F115A* Rev | CCACTATCATCAGAATTATCGCC**AGC**GGTTTTTCTCCCACTTATCACACCCTCAGG | AC7981 |
| *RRP40_164* gRNA Fwd | GCAGTGAAAGATAAATGATC**ACTAGAATAGTCTATATATG**GTTTTAGAGCTAGAAATAGC | AC10660 |
| *SUP4t* crRNA Rev | CCCTCACTAAAGGGAACAAAAGCTGGGTACCAGACATAAAAAACAAAAAAAGCACCACC | AC10140 |
| *rrp40-Y64N* HDR Fwd | AAGAGTGGTGTTCAGAC**T**GCATATATAGACTATTCTAGTAAGAGA**AAC**ATTCCATCTGTAAACGATTTTGTAATCGGTGTCATTATAGGG | AC10661 |
| *rrp40-Y64N* HDR Rev | CCCTATAATGACACCGATTACAAAATCGTTTACAGATGGAAT**GTT**TCTCTTACTAGAATAGTCTATATATGC**A**GTCTGAACACCACTCTT | AC10662 |
| *MPP6-natMX4* Fwd | AAAGTAGTAGAGATGAGGGCCACTTAAACACAATAAGCATATAAGGAACCAAGGGAAAAGCAACAAACATGGAGGCCCAGAATACCCTCC | AC7967 |
| *MPP6-natMX4* Rev | CACAATGCTTTATATTATGTTCTCATTATTATATACGAATACGTACTTCTGCTGGTGCATTGTCTGCGCAGTATAGCGACCAGCATTCAC | AC7968 |
| *U4* pre-snRNA Fwd | AAAGAATGAATATCGGTAATG | AC5722 |
| *U4* pre-snRNA Rev | ATCCTTATGCACGGGAAATACG | AC5723 |
| *TLC1* pre-RNA Fwd | GTATTGTAGAAATCGCGCGTAC | AC7593 |
| *TLC1* pre-RNA Rev | CCGCCTATCCTCGTCATGAAC | AC7594 |
| *U14* snoRNA (snR128) Fwd | GATCACGGTGATGAAAGACTGG | AC5397 |
| *U14* snoRNA (snR128) Rev | CTACAGTATACGATCACTCAGACATCCTA | AC5398 |
| *ALG9* mRNA Fwd | CACGGATAGTGGCTTTGGTGAACAATTAC | AC5067 |
| *ALG9* mRNA Rev | TATGATTATCTGGCAGCAGGAAAGAACTTGGG | AC5068 |
| 7S (ITS2) rRNA Probe | GGCCAGCAATTTCAAGTTA | HG242 |
| 5S rRNA Probe | CTACTCGGTCAGGCTC | HG246 |

**Table S4. MolProbity results for budding yeast and human RNA exosome models**

| **Validation** | **Budding Yeast** | **Human** |
|---|---|---|
| MolProbity score | 1.05 | 1.03 |
| MolProbity Clashscore | 0 | 0 |
| Rotamers outliers (%) | 0.71 | 0.23 |
| Cβ deviations | 0 | 3 |
| Ramachandran favored (%) | 90.02 | 90.51 |
| Ramachandran allowed (%) | 9.20 | 8.65 |
| Ramachandran outliers (%) | 0.78 | 0.84 |
